# Recharacterization of RSL3 reveals that the selenoproteome is a druggable target in colorectal cancer

**DOI:** 10.1101/2024.03.29.587381

**Authors:** Stephen L. DeAngelo, Liang Zhao, Sofia Dziechciarz, Myungsun Shin, Sumeet Solanki, Andrii Balia, Marwa O El-Derany, Cristina Castillo, Yao Qin, Nupur K. Das, Hannah Noelle Bell, Joao A. Paulo, Yuezhong Zhang, Nicholas J. Rossiter, Elizabeth C. McCulla, Jianping He, Indrani Talukder, Billy Wai-Lung Ng, Zachary T. Schafer, Nouri Neamati, Joseph D. Mancias, Markos Koutmos, Yatrik M. Shah

## Abstract

Ferroptosis is a non-apoptotic form of cell death resulting from the iron-dependent accumulation of lipid peroxides. Colorectal cancer (CRC) cells accumulate high levels of intracellular iron and reactive oxygen species (ROS) and are thus particularly sensitive to ferroptosis. The compound (S)-RSL3 ([1S,3R]-RSL3) is a commonly used ferroptosis inducing compound that is currently characterized as a selective inhibitor of the selenocysteine containing enzyme (selenoprotein) Gluathione Peroxidase 4 (GPx4), an enzyme that utilizes glutathione to directly detoxify lipid peroxides. However, through chemical controls utilizing the (R) stereoisomer of RSL3 ([1R,3R]-RSL3) that does not bind GPx4, combined with inducible genetic knockdowns of GPx4 in CRC cell lines, we revealed that GPx4 dependency does not always align with (S)-RSL3 sensitivity, questioning the current characterization of GPx4 as the central regulator of ferroptosis. Utilizing affinity pull-down mass spectrometry with chemically modified (S)-RSL3 probes we discovered that the effects of (S)-RSL3 extend far beyond GPx4 inhibition, revealing that (S)-RSL3 is a broad and non-selective inhibitor of selenoproteins. To further investigate the therapeutic potential of broadly disrupting the selenoproteome as a therapeutic strategy in CRC, we employed additional chemical and genetic approaches. We found that the selenoprotein inhibitor auranofin, an FDA approved gold-salt, chemically induced oxidative cell death and ferroptosis in both *in-vitro* and *in-vivo* models of CRC. Consistent with these data, we found that AlkBH8, a tRNA-selenocysteine methyltransferase required for the translation of selenoproteins, is essential for the *in-vitro* growth and xenograft survival of CRC cell lines. In summary, these findings recharacterize the mechanism of action of the most commonly used ferroptosis inducing molecule, (S)-RSL3, and reveal that broad inhibition of selenoproteins is a promising novel therapeutic angle for the treatment of CRC.

## Introduction

Colorectal Cancer (CRC) is the fourth most common cancer and fifth leading cause of cancer-related deaths in the US^1^. Most early CRC is diagnosed through routine colonoscopies and if found early, CRC is curable with surgical resection and adjuvant chemotherapy^2^. However, recent data demonstrate a concerning rise in the incidence rate of CRC in young adults who often present with advanced disease^3^. Metastatic CRC is incurable and is commonly driven by inactivating mutations of the tumor suppressor genes APC and TP53, leaving minimal opportunities for the development of targeted therapeutics^4,5^. While recent advances in immunotherapy have demonstrated remarkable success in mismatch repair deficient (MMR) CRC^6^, only a small subset of CRC patients are diagnosed with MMR CRC^7^, limiting its widespread utility. Therefore, new effective therapies for CRC are desperately needed.

CRCs are highly addicted to iron^8–11^ and thus an emerging strategy for the treatment of CRC is the induction of an iron-dependent, non-apoptotic form of cell death, ferroptosis^12^. In the presence of labile iron, intracellularly produced hydrogen peroxide undergoes radical decomposition to produce a hydroxyl radical in a process known as the Fenton reaction^13^. The hydroxyl radical rapidly proliferates through cells, leading to widespread DNA, protein, and lipid peroxidation. The accumulation of lipid peroxides is a key characteristic of ferroptosis and results in membrane instability leading to cell rupture^14^. However, in contrast to apoptosis, ferroptosis lacks distinct, well-defined markers for its characterization^15^. Therefore, the main distinguishing characteristic of ferroptosis is its ability to be rescued by the lipid reactive oxygen species (ROS) scavenging antioxidants liproxstatin-1 and ferrostatin-1^16^.

Glutathione peroxidase (GPx)4 utilizes reduced glutathione (GSH) to directly detoxify lipid ROS, leading to the characterization of GPx4 as an essential regulator of ferroptosis^17^. Central to the activity of GPx4 is a catalytic selenocysteine residue as GPx4 is a selenoprotein, one of 25 proteins in the human genome that selectively incorporate the amino acid selenocysteine^18–20^, collectively known as the selenoproteome. The selenocysteine residue is often essential for catalytic function as selenocysteine to cysteine point mutations in GPx4 result in >90% loss of activity, albeit supraphysiological overexpression of this reduced activity mutant rescues ferroptosis induction *in-vitro*^21^. The discovery of <u>R</u>as-<u>S</u>ensitive <u>L</u>igand <u>3</u> (RSL3) – identified as a GPx4 inhibitor^17^ – has further cemented the characterization of GPx4 as the central regulator of ferroptosis^22^. While there is broad cell line sensitivity to RSL3, this does not always correlate with GPx4 expression, casting doubt on RSL3 as a specific inhibitor of GPx4.

Through the generation of doxycycline-inducible GPx4 knockdown cell lines (GPx4 i-KD), we identified a CRC cell line (DLD1) that is sensitive to RSL3 but insensitive to GPx4 i-KD. Next, through synthesis of a biotinylated RSL3, we performed streptavidin-based affinity pulldown of CRC lysates to identify RSL3 targets by tandem-mass tag (TMT) quantitative proteomics. Our data revealed that RSL3 has numerous targets and is a non-specific inhibitor of the selenoproteome and thioredoxin peroxidases, including the selenoprotein Thioredoxin Reductase 1 (TxnRD1). Last, we employed chemical and genetic approaches to modulate the selenoproteome and characterized the effects of selenoproteome inhibition on CRC cell lines. In our studies we demonstrated that the selenoproteome is essential for the growth and survival of CRC cell lines in *in-vitro* and *in-vivo* models, thus revealing the broad inhibition of selenoproteins to be a novel therapeutic strategy in CRC.

## Methods

### Mice

All mice used in these studies are from the Balb/c, C57Bl/6J, or NOD SCID lines as indicated. Males and females are equally represented, and littermates were randomly mixed in all experimental conditions. The mice were housed in a temperature controlled, specific pathogen free environment, with a 12-hour light/dark cycle. They were fed *ad-libitum* with either a standard chow diet or auranofin dosed diet as indicated. Mice were between 6 and 8 weeks old at study initiation. All animal studies were carried out in accordance with the Association for Assessment and Accreditation of Laboratory Animal Care International guidelines and approved by the University Committee on the Use and Care of Animals at the University of Michigan.

### Xenograft studies

CT26 cells were maintained for 2 weeks at < 70% confluency prior to injection into fully immune-competent Balb/c mice. Prior to injection, cells were trypsinized, thoroughly washed, and resuspended in sterile saline to a final injection volume of 100 μL. Wild-type mice of both sexes were anesthetized via inhaled isoflurane (induction dose 3-4%, maintenance dose 1-2%) and inoculated with 0.4 x 10^6^ CT26 cells. Cells were implanted into lower flanks and treatment began on day 4 once palpable tumors were identified. Tumor size was measured with digital calipers utilizing the formula V=0.5*L*W^2^. Xenograft treatments used in this study included the following: auranofin injection (10mg/kg, IP, daily), auranofin chow (1.25-10mg/kg, *ad libitum*), doxycycline chow (400 mg/kg, *ad libitum*), NAC drinking water (20mM, *ad libitum*)

Ethical endpoints for xenograft studies were determined in accordance with Association for Assessment and Accreditation of Laboratory Animal Care International guidelines and approved by the University Committee on the Use and Care of Animals at the University of Michigan. Ethical endpoints for this study were determined to be any of the following: 30 days post implantation, tumor size exceeding 2 cm in any direction, hemorrhage of the tumor, or indications of tumor induced pain and decreased mobility. Once endpoint was determined for any of the groups, all mice in study were euthanized by CO_2_ inhalation and tumors were excised. Tumor volume and weight were measured, and tissues were prepared for histology, IHC, or flow cytometry as indicated. A licensed veterinarian was available for consult at all points during the study.

### AOM/DSS induction of colitis-associated colorectal cancer

C57BL/6J mice were utilized to model colitis-associated colon cancer as previously described^10,23,24^.

Briefly, at 6 weeks of age mice were weighed and injected intraperitoneally with 10 mg/kg of Azoxymethane (AOM). Three days following AOM injection mice began their first cycle of 2% (w/v) Dextran Sodium Sulfate (DSS) water. Following a DSS cycle, mice began a 14-day recovery period before the next cycle of DSS treatment. Treatment with Auranofin chow began 7 days following the conclusion of the 3^rd^ administration of DSS water and continued for 30 days. At the conclusion of the study all mice were euthanized, and the colons were extracted for analysis. Histological analysis was performed via a study-blinded independent pathologist.

### Human Survival Analysis

Human survival analysis was determined using the online tool KMplot^25^. This is a web-based, registration-free survival analysis tool that can perform univariate and multivariate survival analysis. Significance is computed using the Cox-Mantel log-rank test.

### Human Normal vs Tumor Gene Expression Analysis

Human normal vs. tumor gene expression analysis was determined using the online tool UALCAN (University of Alabama at Birmingham Cancer Data Analysis Portal)^26,27^. This is a web-based, registration-free gene expression analysis tool for the analysis of cancer OMICS data from the TCGA (The Cancer Genome Atlas Project).

### Cell lines

Human CRC cell lines HCT116, RKO, SW480, and DLD1, and the mouse CRC cell line CT26 were used. Cell lines have been STR-authenticated and routinely tested for mycoplasma contamination by PCR. All cells were maintained in complete DMEM medium (supplemented with 10% fetal bovine serum (Cytiva) and 1% antibiotic/antimycotic agent (Thermo Fisher)) at 37 °C in 5% CO_2_ and 21% O_2_. Cell numbers were quantified for plating and xenograft experiments using a Multisizer 4e Coulter Counter (Beckman-Coulter).

### Growth assays

Adherent cell growth assays were performed via label free live cell imaging using the Cytation 5 Imaging Multi-Mode reader with attached BioSpa (Agilent BioTek). For 72 hr growth analysis cells were plated at ∼500 cells/well while for 144 hr growth analysis cells were plated at ∼100 cells/well. Cells were allowed to adhere overnight, imaged and analyzed for cell number at t_0_, and then immediately treated as indicated in the figure legend; images were then acquired every 8-24 hr as indicated. Cytation software was used to quantify adherent cell counts.

Analysis was performed by normalization to cell number at first reading (0 hr), with a minimum of 3 independent wells averaged for statistical analysis. Graphs were plotted using Prism with error bars representing mean +/- standard deviation. Growth assay data was utilized for the generation and calculation of EC_50_ curves. The average fold change at the endpoint of an untreated control group (minimum of 3 replicates) was normalized to a value of 1.0 and the relative fold change of treated wells (minimum 3 replicates per group) was then calculated as a fractional response as compared to the average of the untreated control. EC_50_ values were determined using Prism sigmoidal standard curve interpolation. Statistical significance was determined using Prism 2-way ANOVA with multiple comparisons.

### Clonogenic Assays (Colony Formation Assays)

Cells were plated in biological triplicates in a 6-well plate at 500 cells per well in 3 mL of media. Cells were treated as indicated in the figure legends with media and treatment replenished every 5 days. Assays were concluded at 12-15 days by fixing cells in cold 10% buffered formalin for 10 min and staining with 1% crystal violet, and 10% methanol solution for 30 min. Colony plates were washed in dH_2_O and imaged using an iBright FL1500 imaging system (Thermo Fisher)

### Histological Staining

Colonic tissues or tumor tissues were rolled and fixed with PBS-buffered formalin for 24 hours then transferred to 70% ethanol, followed by embedding in paraffin. Sections of 5 μm were stained for H&E and mounted with Permount Mounting Medium (Thermo Fisher Scientific).

### Western Blotting

Indicated cells were seeded in a 6-well plate in triplicate for each condition and allowed to adhere overnight. Cells were plated to reach ∼0.8x10^6^ cells/well at time of harvest. Cells were lysed with RIPA assay buffer with added protease (1:100 dilution; MilliporeSigma) and phosphatase (1:100 dilution; Thermo Fisher Scientific) inhibitors. Lysates were quantified by BCA protein assay kit (Pierce – Thermo Fisher Scientific) and normalized for loading. Solubilized proteins were resolved on 10% SDS-polyacrylamide gels and transferred to nitrocellulose membrane, blocked with 5% milk in TBST, and immunoblotted with the indicated primary antibodies: GPx4 monoclonal (Proteintech 67763-1-Ig), TxnRD1 (Santa Cruz sc-28321), Actin (Proteintech 66009-1-Ig). HRP-conjugated secondary antibodies used were anti-rabbit and anti-mouse at a dilution of 1:2000 and immunoblots were developed using Chemidoc imaging system (ChemiDoc, BioRad).

### CETSA Assay

293T cells cultured in 10 cm dishes were treated with 10 μM compound (1R,3R-RSL3, 1S,3R-RSL3, Auranofin) or DMSO control (0.1%, v/v) for 1 hour at 37 °C. After treatment, cells were harvested and the cell suspensions in PBS were distributed into 12 PCR tubes and heated to indicated temperatures (43.0, 43.5, 45.7, 48.9, 52.4, 55.9, 57.4, 60.5, 63.3, 65.6, 66.3, and 66.7°C) for 3 minutes. After cooled to room temperature for an additional 3 min, cells were lysed by liquid nitrogen with freeze and thaw cycles. Samples were supplemented with protease inhibitor cocktail (#HY-K0010, MCE). Subsequently, the cell lysates were centrifuged at 20,000 g at 4 °C for 20 minutes. The supernatant was carefully collected and diluted with 5× SDS loading buffer (#AIWB0025, Affinibody) for SDS–PAGE (#ET15420LGel, ACE) and western blotting analysis.

### qPCR

Cell lines were treated for indicated times (typically 72 hrs) and washed with sterile PBS prior to RNA extraction with Trizol reagent. Total RNA was visualized on an agarose gel to confirm high quality extraction, with RNA yield quantified using a Nanodrop. 1μg of total RNA was reverse transcribed to cDNA using SuperScriptTM III First-Strand Synthesis System (Invitrogen). Real time PCR reactions were set up in three technical replicates for each sample. cDNA gene specific primers and SYBR green master mix were combined and then run in QuantStudio 5 Real-Time PCR System (Applied BioSystems). The fold-change of the genes were calculated using the ΔΔCt method using *Actb* as the housekeeping mRNA. qPCR primers used: Actin F: CACCATTGGCAATGAGCGGTTC. Actin R: AGGTCTTTGCGGATGTCCACGT AlkBH8 F: AGGTCTTTGCGGATGTCCACGT AlkBH8 R: GAGAGCATCCACCAGTCCACAT

### Generation of Doxycycline Inducible Cell Lines

Doxycycline inducible cell lines were generated using the pLKO.1-Tet On system^28^. shRNA sequences were cloned into the pLKO.1-Tet On backbone, sequence validated via Sanger Sequencing (Genscript), and utilized for lentivirus production through the University of Michigan Vector Core. Cells were transfected with lentiviral particles through spinfection at 900xg, 37 °C, 1 hr in the presence of polybrene (final concentration = 10 μg/mL) then incubated at 37 °C, 5% CO2 for 24 hrs. The next day media was changed and cells were allowed to recover for 24 hrs before addition of puromycin (2 μg/mL) to the culture media. Cells were maintained in puromycin until an untreated control well demonstrated 100% cell death. shRNA sequences: GPx4 sh2: GTGGATGAAGATCCAACCCAA

GPx4 sh3: GCACATGGTTAACCTGGACAA

AlkBH8 sh3: TTACCTGAACACATCATATAT

AlkBH8 sh4: CAGGTGGGAAGGCACTCATTT

### Streptavidin-Affinity Pull Down

Streptavidin-Affinity MS was performed on isolated cell lysates. Cells were plated in 15 cm dishes in normal DMEM (10% FBS, 1% Anti-Anti) and allowed to adhere for 24 hrs. The next day cells were treated with 1 μM selenium in the form of Sodium selenite to increase translation of trace selenoproteins that may otherwise be vulnerable to dropout. Sodium selenite was prepared as a 100 mM stock in ddH_2_O. 24 hrs after administration of selenium, cells were washed with sterile PBS and scraped directly into ice cold RIPA buffer. The protocol was optimized to utilize 10 mg of total lysate per biological replicate to obtain signal over the noise threshold. Soluble lysate was treated overnight at 4 °C with the EC_50_ concentration of the respective compound. The next day Streptavidin-coated magnetic beads (Vector Laboratories) were added and the mixture was incubated with gentle rocking at 4 °C for 2 hrs. Beads were then washed 3x in RIPA using a magnetic separation rack, and a further 3x in SDS-free wash buffer (50 mM Tris-HCl, pH 7.4, 150 mM NaCl, 1 mM TCEP). Beads were then pelleted via centrifugation, flash frozen in LN_2_, and stored at -80 °C prior to MS processing.

### LC-MS/MS

All samples were resuspended in 100 mM 4-(2-Hydroxyethyl)-1-piperazinepropanesulfonic acid (EPPS) buffer, pH 8.5, and digested at 37 °C with trypsin overnight. The samples were labeled with TMT Pro and quenched with hydroxylamine. Samples were desalted via StageTip and dried with speedvac. Samples were resuspended in 5% formic acid, and 5% acetonitrile for LC-MS/MS analysis. Mass spectrometry data were collected using an Astral mass spectrometer (Thermo Fisher Scientific) coupled with a Vanquish Neo liquid chromatograph (Thermo Fisher Scientific) with a 75 min gradient and Nano capillary column (100 µm D) packed with ∼35 cm of Accucore C18 resin (Thermo Fisher Scientific). A FAIMSPro (Thermo Fisher Scientific) was utilized with -30,-35,-45,-55,-60, and -70V for field asymmetric waveform ion mobility spectrometry (FAIMS) ion separations. Data acquisition was performed with a mass range of m/z 350-1350 using a TopSpeed method of 1 s. MS1 resolution was set at 60,000 and singly-charged ions were not sequenced. MS1 AGC target was set as standard, and the maximum ion time was set at 50 ms. For MS2 analysis in the Astral analyzer, only multi-charge state ions (z=2-5) were isolated and fragmented using an HCD collision energy of 35%, an isolation window of 0.5 Th, with a dynamic exclusion of 15 s. MS2 AGC target was set as standard, and the maximum ion time was set at 20ms.

Raw files were searched using the Comet algorithm with a custom database search engine reported previously^29^. Database searching included human (*Homo Sapiens*) entries from UniProt (http://www.uniprot.org, downloaded 2021) with the reversed sequences, and common contaminants (i.e., keratins, trypsin). Peptides were searched using the following parameters: 50 ppm precursor mass tolerance; up to 2 missed cleavages; variable modifications: oxidation of methionine (+15.9949); static modifications: TMTpro (+304.2071) on lysine and peptide N-terminus, carboxyamidomethylation (+57.0215) on cysteines and selenocysteines. The protein-level FDR was determined using the ModScore algorithm where a score of 13 corresponds to 95% confidence in correct localization. TMT reporter ions were used for quantification of peptide abundance. Isotopic impurities were corrected according to the manufacturer’s specifications, and signal-to-noise (S/N) was calculated. Peptides with summed S/N lower than 100 across all channels or isolation specificity lower than 0.5 were discarded. The high confidence peptides were then used to quantify protein abundance by summing up S/N values for all peptides assigned to the same protein, and only proteins in the linear quantification range of the instrument were included in the analysis. The normalization for protein quantification was then performed by adjusting protein loadings of total sum of S/N values to that of every TMT channel.

### Statistical Overrepresentation Test

Statistical overrepresentation analysis/test of proteins pulled down with Biotin-(1S,3R)-RSL3 was performed on the portion of the hit list that met all the following criteria: 1) >1 peptide identified by TMT-MS, 2) p-value < 0.050, 3) fold enrichment > 2. This list of genes was then input into PANTHER database (v18.0) and analyzed for protein class overrepresentation. As selenoproteins are not annotated as a protein class in this current version of PANTHER, we calculated the probability of observing a selenoprotein in a random set of genes from the human genome as 25/20592 (utilizing the reference list size of PANTHER v18.0). This probability value was scaled to our input list and a Fisher’s exact t-test was performed for statistical significance.

### C11-BODIPY lipid ROS measurement

Indicated cells were seeded in 12-well plates and allowed to adhere overnight at 37 °C prior to beginning the indicated treatment or targeted gene knockdown induced by doxycycline. Cells were harvested using PBS-EDTA (5 mM), buffer, washed once with HBSS, suspended in HBSS containing 5 μM C11-BODIPY (Thermo Fisher), and incubated at 37 °C for 30 min. Cells were pelleted, washed, and resuspended in HBSS. Fluorescence intensity was measured on the FITC channel using the Beckman Coulter MoFlo Astrios. A minimum of 20,000 cells were analyzed per condition. Data were analyzed using FlowJo software (Tree Star). Values are expressed as MFI.

### Synergy Calculations

Synergy was calculated through disproof of the null hypothesis that treatment with two biologically active compounds would produce a result in line with the Bliss Model of Independence as previously described^30^.

### ICP-MS

Whole blood from indicated mice were obtained via submandibular vein puncture or from the orbital sinus. Whole blood was collected in untreated sterile 1.5 mL Eppendorf tubes and allowed to coagulate for 1-2 hrs at RT. Coagulated samples were spun at 13,000xg for 15 min and serum was collected. Collected serum was further clarified with a second spin at 13,000xg for 15min. 10 μL of clarified serum was treated with 2 mL/g total wet weight nitric acid (20 μL) (Trace metal grade; Fisher) for 24 hr, and then digested with 1 mL/g total wet weight hydrogen peroxide (10 μL) (Trace metal grade; Fisher) for 24 h at room temperature. The samples were preserved at 4 °C until quantification of metals. Ultrapure water (VWR Chemicals ARISTAR®ULTRA) was used for final sample dilution to 3 mL. Samples were then analyzed using inductively coupled plasma mass spectrometry (ICP-MS) (Perkin Elmer) utilizing Bismuth as an internal standard.

### Auranofin Diet

Auranofin diet was made from powdered laboratory rodent diet (LabDiet) mixed with appropriate quantities of drug in a KitchenAid mixer designated for laboratory use. Diet dosing was calculating assuming an average mouse weight of 25 g and chow consumption of 4 g/day as previously published^31^. Water was added to a hydration level of 60% (600 mL H_2_O per kg of diet) and thoroughly mixed. Small quantities of food-grade coloring (McCormick) were added to differentiate doses of diet. Following mixing, the diet was extruded into pellets and dehydrated for 72 hr with a food dehydrator at 41 °C (Nesco). Mice were not provided an alternate food source when undergoing treatment and weight was routinely monitored.

### Thioredoxin Reductase Activity Assay

Thioredoxin Reductase (TxnRD) activity assay was based on the plate reader procedure as outlined by Cunnif *et al.*, Anal. Biochem, 2013^32^ and established by Arnèr *et al*., Methods in Enzymology, 1999^33^. Briefly, the TxnRD activity assay is based on consumption of the diselenide amino acid selenocystine in cell lysate, measured by NADPH consumption via absorbance at 340 nm on a spectrophotometric plate reader (Cytation 5, BioTek). TxnRD is the only cellular enzyme capable of reducing selenocystine, consuming NADPH in the process. A master-mix of 2 mM NADPH and 1 mM selenocystine was prepared fresh for each assay. NADPH (Sigma Aldrich) solution was prepared in 100 mM Tris, pH 8; fresh solution was used whenever possible and solution was used within 1 week if frozen at -20 °C, with reduced initial absorbance observed when utilizing thawed NADPH. As selenocystine does not readily dissolve in aqueous buffer, selenocystine (Cayman Chemical Company) was first dissolved in 1N NaOH (650 μL for a 50 mg vial), to which ½ volume of 1N HCl was added to neutralize the solution. The master stock was then diluted to 45 mM in ddH2O and stored at -20 °C. Cells were plated and treated as indicated, washed in PBS, and lysed in RIPA buffer. The insoluble fraction was removed via centrifugation (13,000xg, 10 min, 4 °C) and protein abundance was quantified by BCA Protein Assay (Thermo Scientific). 50 μg of protein was utilized per well in a total volume of 60 μL, to which 40 μL of NADPH/Selenocystine master-mix was added. Immediately following addition of the master-mix (within 2 min), kinetic reading of absorbance values was initiated, with reads taken every 60 s for 30 min.

### CRISPR Co-Essentiality Network Generation

CRISPR gene effect scores from the DepMap 22Q2 release were first corrected using Cholesky whitening as previously described^34^. A matrix of p-values corresponding to gene-wise Pearson correlations was calculated from the whitened data, then converted to FDR values using the p.adjust() function in R. A network was then constructed from all partners within 2 edge distances from a given gene of interest, using an FDR cutoff of 0.05 to define edges/partnership. Network graphs were generated from pairwise gene lists using the tidygraph and ggraph packages in R, with the layout argument in ggraph set to “graphopt”

### AGB 364/366 synthesis

**Scheme 1.**
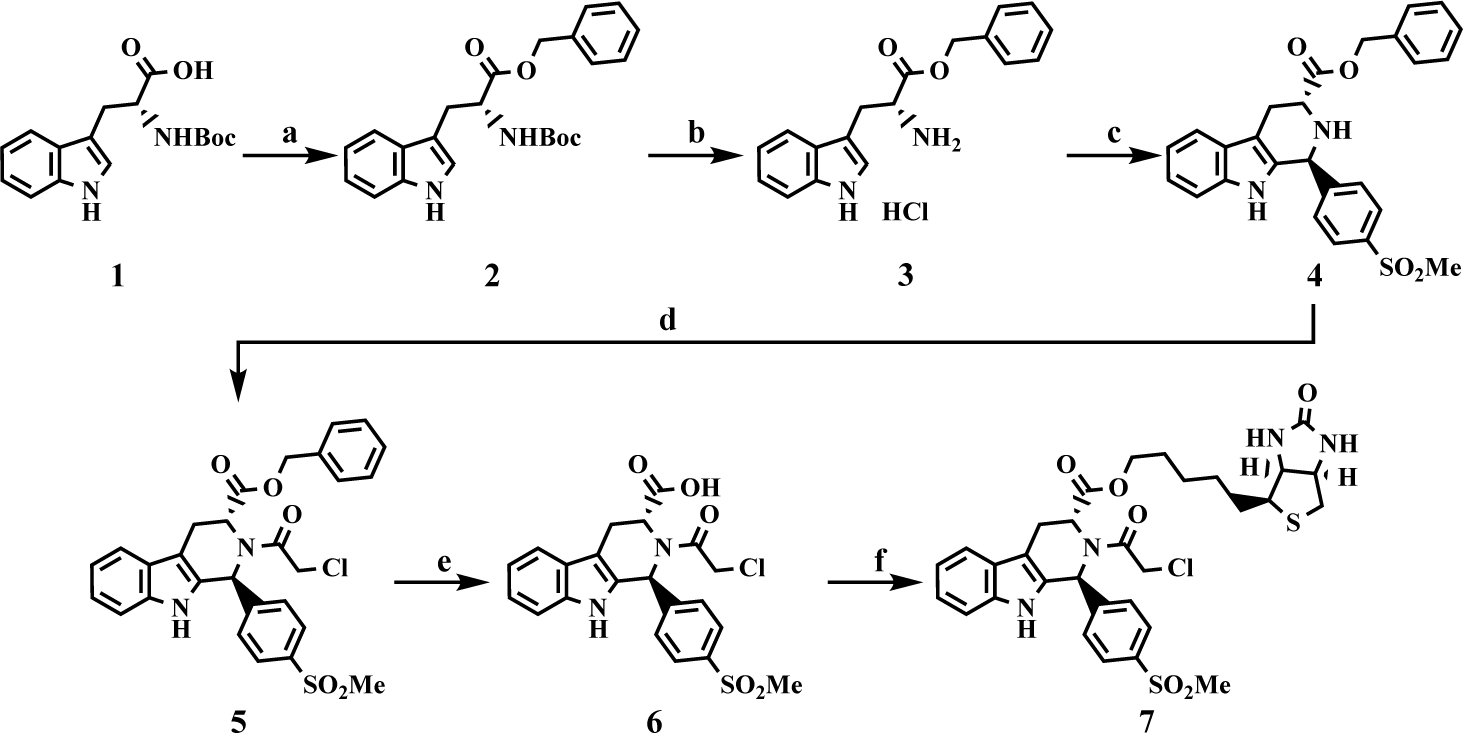
Synthesis of Cpd24 and Biotinylated Analog^a^. ^a^Reagents and conditions: (a) BnOH, EDC, DMAP, THF, rt, overnight; (b) 4 N HCl in 1,4-dioxane, rt, overnight, 92% over 2 steps; (c) 4-(methylsulfonyl)benzaldehyde, IPA, reflux, 87%; (d) ClCH_2_C(O)Cl, TEA, CH_3_CN, reflux, 87%; (e) H_2_, Pd/C, EtOH, rt, overnight, 72%; (f) *D*-biotinol, EDC, DMAP, THF, rt, overnight, 4%.

**Scheme 2.**
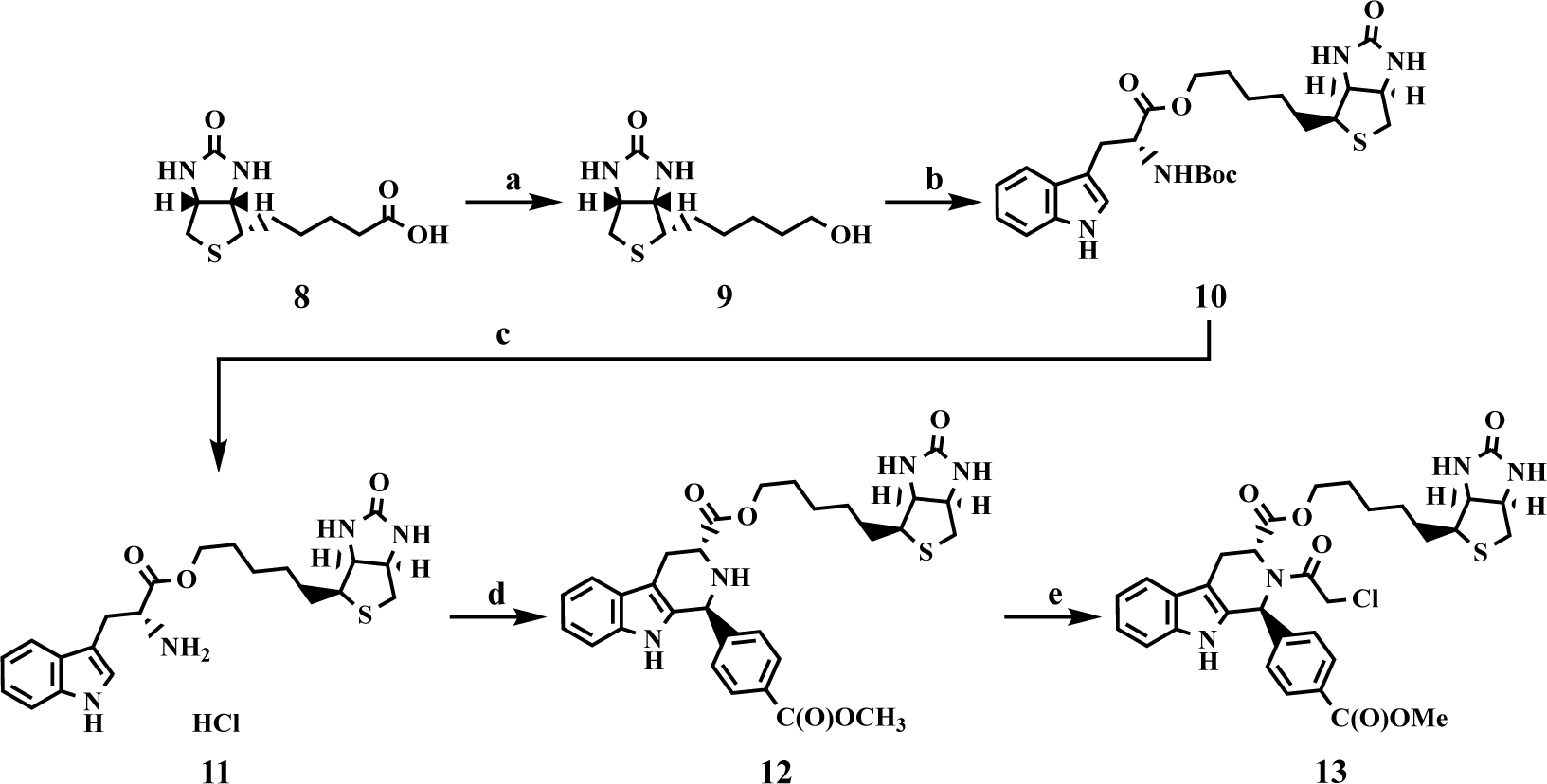
Synthesis of RSL3 and Biotinylated Analog^b^. ^b^Reagents and conditions: (a) LiAlH_4_, THF, reflux, overnight, 45%; (b) (*tert*-butoxycarbonyl)-*D*-tryptophan, EDC, DMAP, THF, rt, overnight, 74%; (c) 4 N HCl in 1,4-dioxane, rt, overnight, 91%; (d) methyl 4-formylbenzoate, IPA, reflux, 15%; (e) ClCH_2_C(O)Cl, TEA, CH_3_CN, reflux, 19%.

### Statistics

Data are represented as mean ± standard deviation. unless otherwise indicated. Data are from a minimum of 3 independent experiments measured in triplicate unless otherwise stated in the figure legend. For statistical analyses, unpaired *t*-tests were conducted to assess the differences between 2 groups. One-way or 2-way ANOVA was used for multiple treatment conditions followed by Tukey’s post hoc test. A strict p-value cutoff of < 0.050 was utilized for determination of significance in all experiments. All statistical tests were carried out using Prism 10 software (GraphPad).

### Study approval

All animal studies were carried out in accordance with Institute of Laboratory Animal Resources guidelines and approved by the University Committee on the Use and Care of Animals at the University of Michigan (IACUC protocol number: PRO00011805)

## Results

### The potency of (S)-RSL3 is due to GPx4 independent effects in CRC cell lines

The most commonly utilized mechanism of ferroptotic induction is treatment with the small molecule GPx4 inhibitor RSL3 (Figure 1A). RSL3 has two stereoisomers, (1S,3R)-RSL3 and (1R,3R)-RSL3, which we will refer to as (S) and (R)-RSL3, respectively (Figure 1B). The (S) and (R) stereoisomers of RSL3 differentially target GPx4^22^ (Figure 1C, S1A), with the ability of (S)-RSL3 to more potently inhibit GPx4 hypothesized to directly correlate with its increased potency. Utilizing 72 hr live cell imaging growth assays we confirmed that (S)-RSL3 is markedly more potent than (R)-RSL3 in the HT1080 fibrosarcoma cell line that is frequently utilized in ferroptosis research^35^. The EC_50_ values of 13 nM for (S)-RSL3 and 903 nM for (R)-RSL3 (Figure 1D-E) are consistent with previously reported cell death and viability measurements^17^. However, both (S)-RSL3 and (R)-RSL3 exhibited nearly equipotent sensitivity in CRC cell lines DLD1, HCT116, RKO, and SW480, with EC_50_ values at ∼1 μM (Figure 1F). Notably, DLD1 cells displayed a 2-fold increased sensitivity to (R)-RSL3, despite the decreased affinity of (R)-RSL3 towards GPx4.

**Figure 1:**
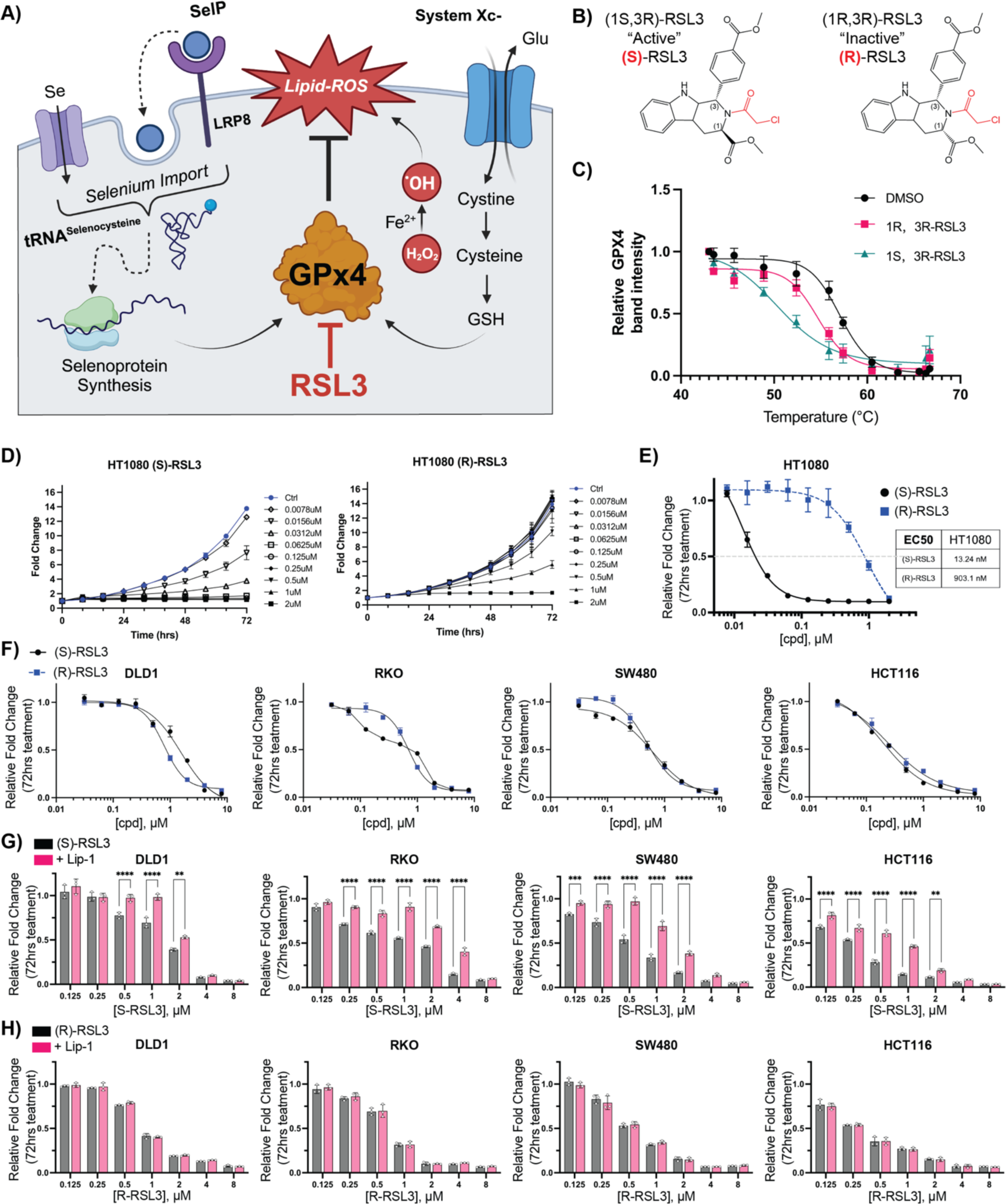
Defining the ferroptotic window of S-RSL3 in CRC cell lines. **A)** Schematic of ferroptosis demonstrating the role of of GPx4 to detoxify ROS induced lipid peroxides (Lipid-ROS) is regulated by GSH synthesis driven by System Xc-import of cystine and selenium availability in part by LRP8 which is then incorporated into the GPx4 polypeptide by tRNA-selenocysteine. **B)** Structures of (1S,3R)-RSL3 and (1R,3R)-RSL3. **C and D)** Cell growth assay normalized to untreated control at 72 hr following (S) or (R) RSL3 treatment in HT1080 and CRC cell lines. **E)** Results of cell growth assay from C-D where 72 hr growth of cells treated with indicated doses of (S) and (R) RSL3 is normalized to 72 hr growth of vehicle treated control wells to calculate EC50 value (“Results of 72 hr cell growth assay”). **F)** Results of 72 hr cell growth assay of (S) and (R) RSL3 across a panel of CRC cell lines. **G-H)** Statistical analysis of results of 72 hr cell growth assay of (S) and (R) RSL3 co-treated with liproxstatin-1 (Lip-1) (1 μM) *: p < 0.05, **: p < 0.01, ***: p < 0.001, ****: p < 0.0001

In contrast to other forms of regulated cell death, ferroptosis lacks specific cellular markers and is instead assessed through the ability to rescue cell death with lipid-ROS scavenging agents such as Liproxstatin-1 (Lip-1)^16^. In our growth assays, we aimed to delineate the “ferroptotic window” induced by (S) and (R)-RSL3 through Lip-1 co-treatment. Despite broad cell line sensitivity to (S)-RSL3, the ferroptotic window varied significantly between cell lines and often did not result in a complete rescue of (S)-RSL3 treatment (Figure 1G, S1B). In agreement with previous data, (R)-RSL3 decreased cell growth, which was not rescued by Lip-1 at any dose in our cell line panel (Figure 1H, S1C). The equipotent nature of these two compounds suggest that while GPx4 inhibition likely drives ferroptosis, the potency of (S)-RSL3 in CRC cell lines may be driven by a GPx4-independent activity.

### RSL3 sensitivity does not predict GPx4 essentiality in CRC cell lines

To investigate if (S)-RSL3 functions independently of its ability to inhibit GPx4 in CRC, we generated stable cell lines with doxycycline-inducible GPx4 shRNAs (GPx4 i-KD). Two RNA guides for GPx4, sh2 and sh3, were found to decrease GPx4 levels by >95% at 72 hr post doxycycline treatment (Figure 2A, S2A). GPx4 was confirmed to be essential for cell growth in the CRC cell lines SW480, RKO, and HCT116, however DLD1 cells were insensitive to GPx4 i-KD despite sensitivity to (S)-RSL3 (Figure 2B). Furthermore, near complete GPx4 i-KD in DLD1 cells resulted in a 10-fold increase in sensitivity to (S)-RSL3 (Figure 2C), supporting the hypothesis that the potency of (S)-RSL3 in CRC cells can be due to GPx4-independent effects. We additionally tested a next generation GPx4 inhibitor, the ML210^36^ derivative JKE1674^37^. JKE1674 is markedly less potent than (S)-RSL3 across CRC cell lines and is inactive up to 20 μM in the DLD1 cell line (Figure 2D). JKE1674 exhibited significantly lower potency in CRC lines as compared to (S)-RSL3, and the decreases in growth induced by JKE1674 treatment up to 20 μM were completely rescued by Lip-1, as demonstrated in the RKO cell line (Figure 2E).

**Figure 2:**
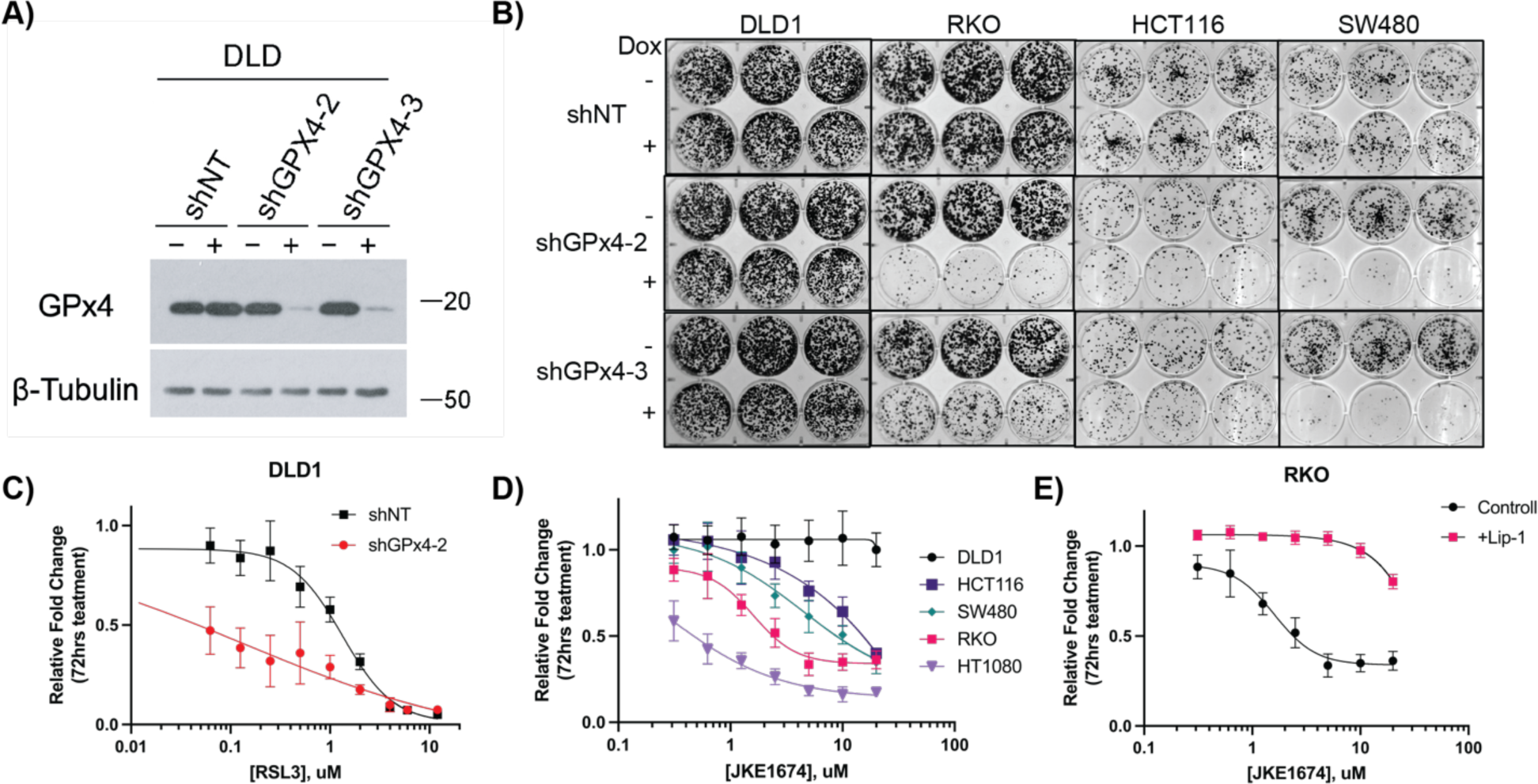
Elucidating the role of GPx4 in CRC cell lines. **A)** Western blot of stable DLD1 doxycycline inducible shRNA cell lines (shNT, shGPx4-2, shGPx4-3), +/- doxycycline treatment (250 ng/mL) assessed at 72 hrs. β-Tubulin was used as a loading control. **B)** Colony formation assays of indicated stable shGPx4 or shNT cell lines treated +/- doxycycline (250ng/mL) assessed at 2 weeks. **C)** Results of 72 hr cell growth assay following (S)-RSL3 treatment in DLD1 shNT vs shGPx4-2 cell lines, pre-treated with doxycycline (250 ng/mL) for 72 hrs prior to addition of (S)-RSL3. **D)** Results of 72 hr cell growth assay following JKE1674 dose response of CRC cell lines and HT1080 cells. **E)** Results of 72 hr cell growth assay following JKE1674 dose response of the RKO CRC cell line +/- Lip-1 (1 μM) co-treatment

Altogether, these data demonstrate that the DLD1 cell line is insensitive to modulation of GPx4 activity despite sensitivity to (S)-RSL3. Recent data have suggested that a target of (S)-RSL3 is the selenoprotein Thioredoxin Reductase (TxnRD1)^38^, which regulates the activity of thioredoxin peroxidases, also known as peroxiredoxins (PRDXs) ^39,40^(Figure S2B). Utilizing an in-house thioredoxin reductase activity assay we confirm that (S)-RSL3 also potently inhibits TxnRD activity. Interestingly, DLD1 cells had the highest TxnRD activity across tested CRC lines (Figure S2C-E).

### Chloroacetamide-based ferroptosis inducers non-selectively inhibit the selenoproteome

To determine the targets of (S)-RSL3 in CRC cells we synthesized a biotinylated derivative, utilizing the same attachment site at the 3 position as the original fluorescein conjugate utilized for chemoproteomics in 2014^17^ (Figure 3A). The synthesis yielded a small quantity of biotinylated (R)-RSL3 which we used to validate that the biotin group did not significantly alter the potency of these compounds (Figure 3B). Following optimization of pull-down protocol, including 1 μM selenium supplementation to increase trace selenoprotein expression (Figure S3A), streptavidin-based pull-downs of (S)-RSL3 vs biotin-(S)-RSL3 were performed on biological triplicates. The use of Tandem Mass Tag (TMT) labeling combined with the use of next-generation mass spectrometers has offered unparalleled sensitivity, revealing that while (S)-RSL3 does inhibit GPx4, it is also a broadly non-specific inhibitor (Figure 3C). However, we suspect that many of these targets may not be pharmacologically relevant, as the chloroacetamide moiety utilized in the RSL3 “warhead” is known to be highly reactive and broadly non-specific^41^. To further explore the targets of chloroacetamide-based ferroptosis inducers we synthesized a biotinylated version of the RSL3 derivative Cpd24^42^, observing near identical results to biotinylated (S)-RSL3 (Figure S3B). Statistical overrepresentation analysis revealed two heavily enriched protein classes as targets of (S)-RSL3, peroxidases and selenoproteins (Figure 3D-E).

**Figure 3:**
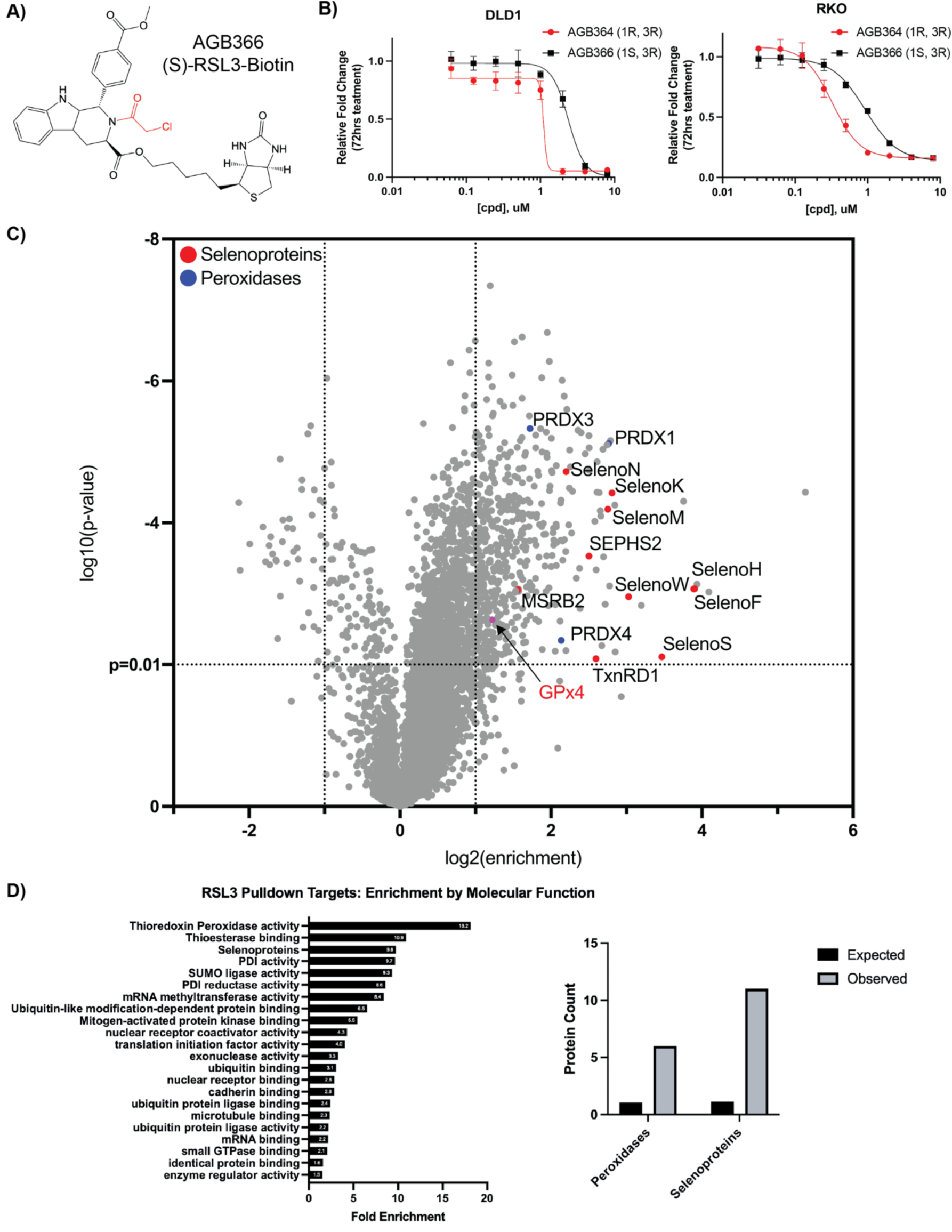
Affinity pulldown-mass spectrometry redefines RSL3 as a selenoprotein inhibitor. **A)** AGB366, a biotinylated RSL3 derivative. **B)** Results of 72 hr cell growth assay following AGB364/366 dose response treatment in DLD1 and RKO cells. **C)** Affinity-pulldown mass spectrometry analysis of AGB366 ((S)-RSL3-Biotin) from RKO lysate with targets of interest identified. **D)** RSL3 Pulldown targets protein class enrichment (PANTHER) and number of peroxidases and selenoproteins observed in the AGB366 pulldown dataset vs. expected by random chance.

### Gold therapy inhibits the selenoproteome and can induce *in-vitro* ferroptosis in CRC

Our data suggests that in addition to its ability to inhibit GPx4 and TxnRD1 – disrupting both cellular mechanisms of peroxide detoxification (Figure 4A) – the potency of (S)-RSL3 may also arise from its ability to broadly inhibit the selenoproteome. We hypothesized that alternative strategies of non-specific selenoprotein inhibition may also be capable of inducing oxidative stress and ferroptosis in CRC cells. To test this hypothesis we used the gold salt containing small molecule auranofin, as ionic gold potently forms Au-Se inhibitory adducts^43,44^. Auranofin is primarily characterized as a TxnRD1 inhibitor^45,46^ with the ability to non-specifically inhibit other selenoproteins^47–49^. However, all small molecules have intrinsic bias in their protein targets and in our hands by CETSA analysis, auranofin showed no statistically significant interaction with GPx4 (Figure S4A-B), providing a unique opportunity to study broad disruption of the selenoproteome without the dominant effects of GPx4 inhibition in these ferroptosis sensitive cell lines.

**Figure 4:**
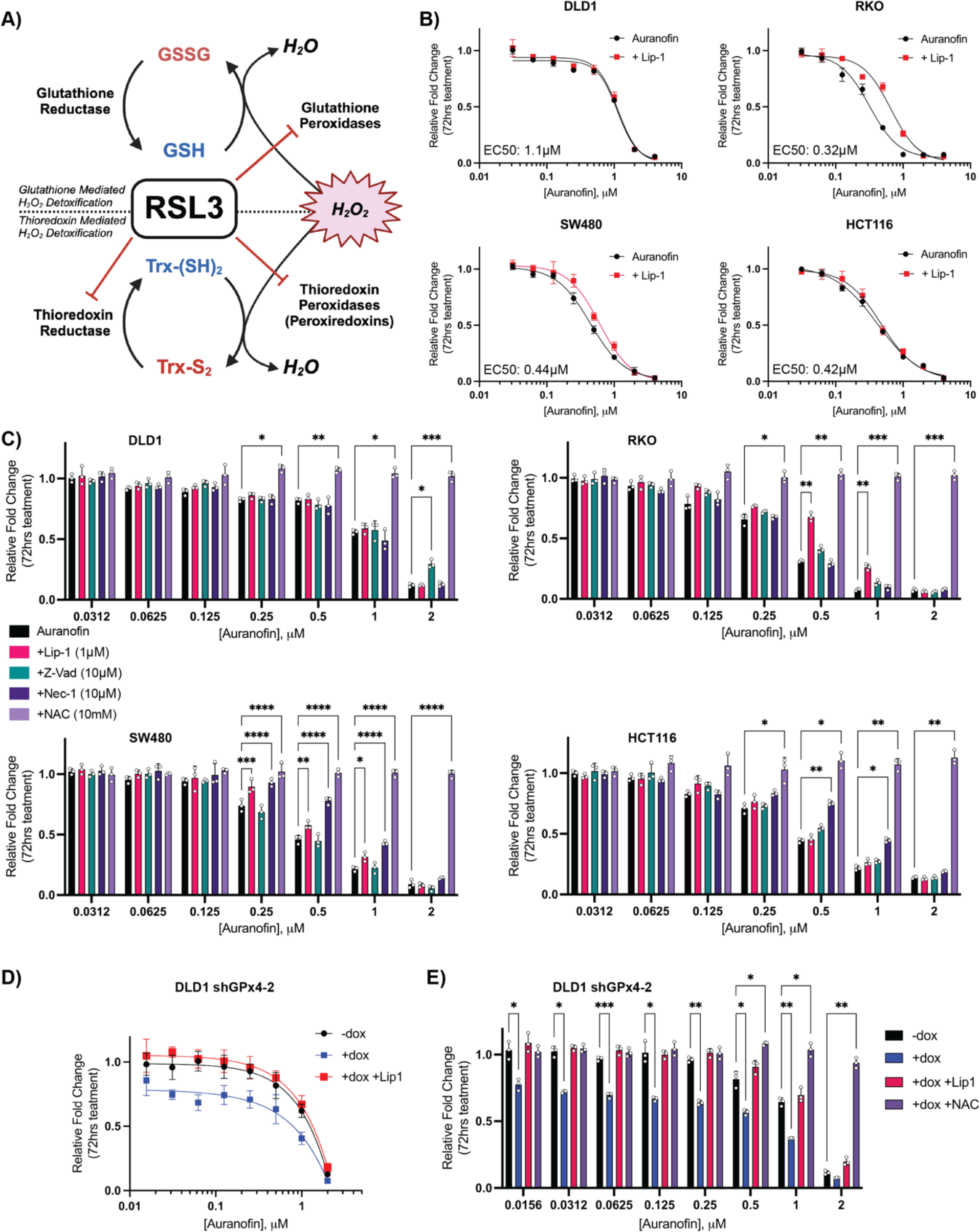
Auranofin induces ferroptosis in CRC. **A)** Schematic of the new proposed mechanism of RSL3 activity where RSL3 as a pan-inhibitor of the selenoproteome can inhibit both glutathione and thioredoxin reductases as well as peroxiredoxins. **B)** Results of 72 hr cell growth assay following auranofin +/- Lip-1 (1 μM) co-treatment across a panel of CRC cells. **C)** Statistical analysis of 72 hr cell growth assay rescue by indicated co-treatments: Lip-1 (1 μM), Z-vad-FMK (10 μM), NAC (10 mM), Nec1 (10 μM). Cells were pre-treated with rescue agent for 24 hr prior to addition of auranofin **D)** Results of 72 hr cell growth assay of the DLD1 shGPx4-2 cell line treated as indicated (vehicle treated control, 250 ng/mL doxycycline, 1 μM Lip-1). Cells were pretreated with doxycycline or vehicle (ddH_2_O) for 72 hr prior to first imaging and treatment with auranofin +/- Lip-1 **E)** Statistical analysis of 72 hr cell growth assay of DLD1 shGPx4-2 cells measuring deviation of growth response following auranofin treatment from control (-dox) by doxycycline treatment (+dox) +/- co-treatment with Lip-1 (1 μM) or NAC (10mM) *: p < 0.05, **: p < 0.01, ***: p < 0.001, ****: p < 0.0001

To investigate the therapeutic potential of alternative selenoprotein inhibition strategies in CRC we initially performed dose response curves across our CRC cell line panel, observing that CRC cells display broad and potent sensitivity to auranofin. Furthermore, several cell lines (RKO, SW480) displayed statistically significant rescue of auranofin treatment by Lip-1 (Figure 4B), demonstrating that gold-based inhibition of the selenoproteome can induce ferroptosis (in a dose and cell line specific manner).

We next sought to further characterize the mechanism of auranofin based growth inhibition, as auranofin is a potent inducer of apoptosis in other cell lines^50^. Supporting a primary role of selenoproteins as antioxidant enzymes, all tested concentrations of auranofin were rescued by the general antioxidant N-Acetyl Cysteine (NAC) while the apoptosis inhibitor Z-Vad-FMK^51,52^ or the necroptosis inhibitor Nec-1^53^ (Figure 4C) induced a slight cell line specific rescue. Therefore, mechanisms of growth inhibition outside of ferroptosis induced by selenoprotein inhibitors appear to also be both dose and cell line dependent. However, when inhibitor concentrations are increased to the μM range, growth can no longer be rescued with any specific cell death inhibitor, supporting a hypothesis that high doses of auranofin induce non-specific oxidative necrosis, consistent with our observations for (S)-RSL3.

To more fully recapitulate the proposed mechanism of (S)-RSL3, the potency of auranofin was tested in combination with GPx4 KD. As we have previously demonstrated, the DLD1 shGPx4-2 cell line is insensitive to GPx4 i-KD, yet combination with GPx4 KD synergistically increased the potency of auranofin treatment (Figure 4F). This combination also drastically shifted the primary mechanism of lower dose auranofin treatment to ferroptosis as determined by Lip-1 rescue (Figure 4E). Altogether these data support a model where (S)-RSL3 activity is driven by its ability to function as a pan-inhibitor of the selenoproteome to broadly inhibit cellular antioxidant systems.

### Gold therapy reduces CRC growth *in-vivo*

We next sought to validate our *in-vitro* findings in an *in-vivo* setting. For these studies, we utilized the CT26 cell line, a mouse derived colorectal adenocarcinoma cell line suitable for growth in fully immunocompetent Balb/c mice. After confirmation that CT26 cells were responsive to auranofin treatment, Lip-1 rescue demonstrated that CT26 cells possess a large and potentially targetable ferroptotic window where doses of auranofin between 60nM and 2μM can be significantly rescued with Lip-1 (Figure 5A). Following flank implantation of 0.4 x 10^6^ CT26 cells per mouse, treatment with auranofin (10 mg/kg, IP, daily) began once tumors were palpable at day 4. At the primary study endpoint (tumor size >2 cm in any direction within a control mouse), IP-treated mice displayed a consistent ∼30% reduction in final tumor mass (Figure 5B), with histological staining demonstrating signs of intra-tumoral necrosis (Figure 5C). However, daily IP of auranofin was not well tolerated as treated mice displayed weight loss and increased stress on handling. As auranofin is orally bioavailable, custom rodent chow was made in-house in a range of concentrations providing dosing from 1.25-10 mg/kg based on an average daily consumption of 4g chow per mouse per day^31^. Additionally, as auranofin rapidly dissociates in plasma to release ionic gold, the bioavailability of auranofin administration can be estimated through analysis of serum gold concentration using inductively coupled plasma mass spectrometry (ICP-MS). Dose optimization studies on non-tumor bearing mice demonstrate that diet administration of auranofin allows for an accurate and reproducible *in-vivo* dose response, with identical endpoint plasma gold concentrations observed between 10 mg/kg administration by chow or IP (Figure S5A). Dose optimization studies also demonstrated that auranofin administration via chow was much better tolerated than IP administration (Figure S5B). Furthermore, ICP-MS data demonstrated that mice administered 10 mg/kg auranofin display plasma gold levels equivalent to patients receiving 6 mg/day auranofin^54,55^ (0.5-0.7 μg/mL), estimated at plasma concentrations equivalent to ∼1-2 μM auranofin^56^.

**Figure 5:**
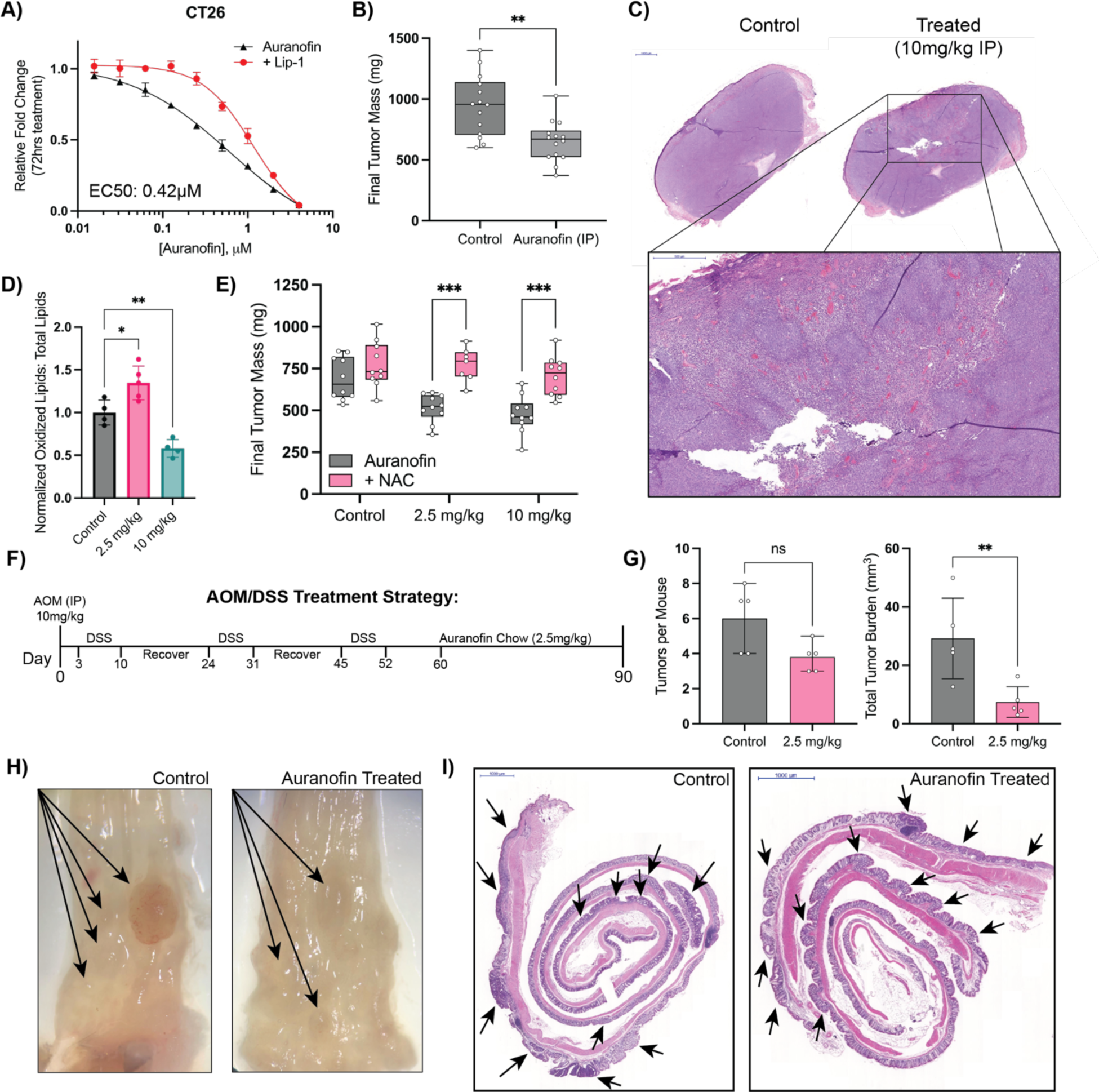
Auranofin is an effective treatment in *in-vivo* models of CRC. **A)** Results of 72 hr cell growth assay of auranofin dose response +/- Lip-1 (1 μM) co-treatment in CT26 cells. **B)** Final tumor mass of CT26 xenograft studies in Balb/c mice treated with vehicle or auranofin IP injection (10mg/kg daily) **C)** H&E staining of Auranofin treated vs control tumor with higher magnification of a region of interest in the treated tumor. **D)** Flow cytometry measurements of Lipid-ROS levels in control vs auranofin treated CT26 xenografts. **E)** Final tumor mass of CT26 xenografts treated with indicated doses of auranofin chow +/- NAC (20mM) drinking water. **F)** Schematic of AOM/DSS model of colitis induced colorectal cancer and subsequent auranofin treatment **G)** Quantification of tumor number and total tumor burden (mm^3^) per mouse colon following AOM/DSS induction and subsequent auranofin treatment. **H)** Representative microscopy pictures of control and treated colons (Auranofin 2.5 mg/kg) with visually detectable tumors marked by arrows. **I)** H&E staining of swiss-rolled colons from C. Notable regions of dysplasia are marked by arrows. *: p < 0.05, **: p < 0.01, ***: p < 0.001, ****: p < 0.0001

We next repeated our CT26 xenograft study with our in-house formulated dose response chow. Doses of 2.5mg/kg and 10mg/kg were chosen to test effects of both low and high dose auranofin in this tumor model. Interestingly, we observed equal effects in tumor growth reduction at both tested doses, which were fully rescuable with co-administration of NAC in drinking water (Figure 5D). Flow cytometry analysis of auranofin-treated tumors confirmed differential *in-vivo* mechanisms, with only low dose auranofin chow capable of increasing intra-tumoral lipid-ROS (Figure 5E).

However, CRC has a unique microenvironment in the colon that is not recapitulated in a flank xenograft model. Therefore, we aimed to use the AOM/DSS model of chemical carcinogen-induced colorectal cancer to study the effects of auranofin on a diverse population of cells within the native microenvironment (Figure 5F). We chose to use 2.5 mg/kg auranofin chow for this study as it was the lowest effective dose in flank xenograft studies. Furthermore, as 2.5 mg/kg auranofin chow was observed to induce lipid-ROS in our flank tumors, this dose allowed us to study the effects of *in-vivo* induction of ferroptosis on CRC tumor progression. Mice were administered a single injection of AOM intraperitoneally followed by three 7-day cycles of DSS with a two-week recovery period between cycles. Following the last cycle of DSS, mice were given a brief recovery period of 7 days and then administered auranofin chow *ad-libitum* until study endpoint. Mice were euthanized after 30 days on auranofin chow. Total number of tumors per mouse trended lower in the treated mice, there was a significant reduction in tumor size and total tumor burden in the auranofin treated group (Figure 5G). While control mice often had large, vascularized colon tumors, the tumors of the auranofin treated group failed to progress (Figure 5H). Histological analysis further confirmed the presence of multiple areas of neoplasia in both groups (Figure 5I), with the tumors of the control group significantly larger than those in the treated group.

### The tRNA-Sec methyltransferase AlkBH8 is required for CRC growth *in-vitro*

To confirm that general inhibition of the selenoproteome is an efficacious target for therapeutic intervention in CRC, a genetic approach to modulate the selenoproteome was required. To guide our approach, co-essentiality analysis revealed that Gpx4 and TxnRD1 are co-essential with the bulk of the tRNA-sec biosynthetic pathway (Figure S6A-B). From these analyses, we identified the tRNA-methyltransferase AlkBH8 as a potential novel therapeutic target in CRC, as increased AlkBH8 expression most significantly correlates with decreased overall survival in CRC patients (Hazard Ratio = 1.6, p = 0.034) as compared to other members of the tRNA-sec biosynthetic pathway (Figure 6A, S6C). AlkBH8 has been extensively reported to regulate the selenoproteome via methylation of tRNA-selenocysteine, with AlkBH8 knockdown and knockout models demonstrating a decreased ability to translate selenoproteins^57–64^. From analysis of TCGA banked tumor samples we observed that AlkBH8 expression is significantly increased in primary tumor samples as compared to normal controls (Figure 6B), and that patients with early-onset CRC have the most highly increased expression of AlkBH8 as compared to other age groups (Figure 6C). As AlkBH8 possesses two druggable domains^60,65^ and could be a target for future development of targeted therapies for early onset-CRC, we sought to experimentally investigate the role of AlkBH8 in regulating the selenoproteome in CRC.

**Figure 6:**
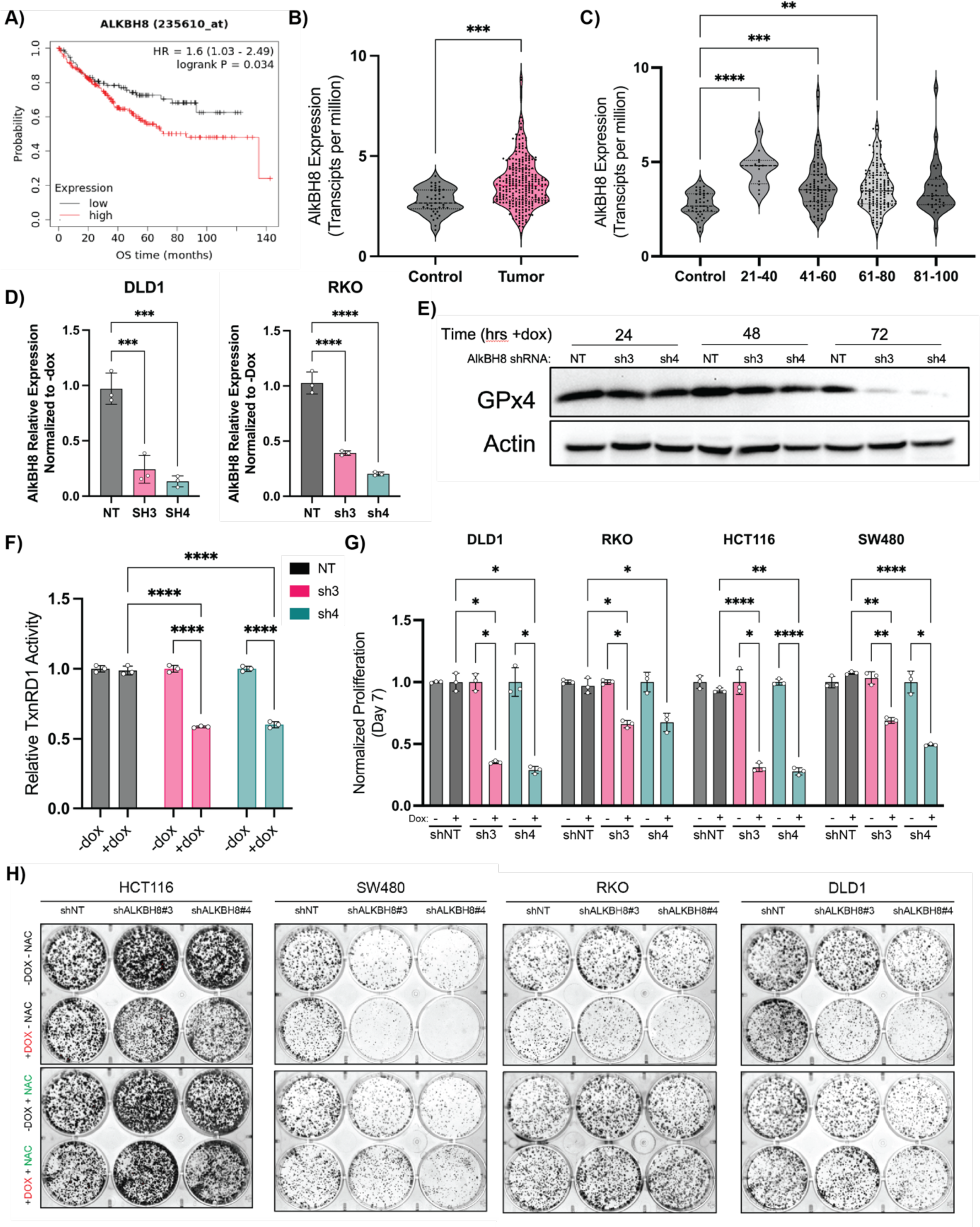
AlkBH8 is a potential therapeutic target in CRC. **A)** Kaplan-Meier overall survival analysis of CRC patients based on AlkBH8 expression as measured by RNA-Seq. **B)** RNA analysis of colorectal adenocarcinoma (COAD) tumor samples vs normal tissue (TCGA) analyzed for AlkBH8 expression. **C)** RNA analysis of colorectal adenocarcinoma (COAD) tumor samples vs normal tissue (TCGA) analyzed for AlkBH8 expression and separated by age group. **D)** qPCR measurement of AlkBH8 mRNA levels in NT and shRNA cell lines following 72 hrs treatment with doxycycline (250ng/mL). Normalized to untreated control. **E)** Western blot analysis of GPx4 protein levels in DLD1 shNT, shAlkBH8-3, and shAlkBH8-4 cell lines following doxycycline treatment (250ng/mL) for indicated times. **F)** TxnRD1 activity in DLD1 shNT, shAlkBH8-3, and shAlkBH8-4 cell lines following doxycycline treatment (250ng/mL) for 72hrs **G)** Cell growth assay at 7 days of indicated CRC cell lines stably transduced with either shNT, shAlkBH8-3, or shAlkBH8-4 and continually treated with doxycycline (250ng/mL). Normalized to untreated control growth at d7. **H)** CFA analysis of CRC cell lines stably transduced with either shNT, shAlkBH8-3, or shAlkBH8-4 as indicated and continually treated +/- doxycycline (250ng/mL) and +/- NAC (10mM) as indicated *: p < 0.05, **: p < 0.01, ***: p < 0.001, ****: p < 0.0001

Two shRNA guides for AlkBH8 i-KD (sh3/4) were found to induce knockdown of AlkBH8 by 60% (sh3) and 80% (sh4) respectively and were cloned into pLKO.1-Tet On for lentivirus based production of stable doxycycline inducible KD cell lines (Figure 6D, S6D). In agreement with prior literature, we observed that AlkBH8 i-KD led to an inability to translate GPx4 as western blots in DLD1 stable shRNA cell lines revealed a sharp drop-off in GPx4 protein concentrations at 72 hrs following doxycycline treatment (250ng/mL) in sh3/4 cell lines but not the NT control line (Figure 6E). Furthermore, we observed a 40% reduction in TxnRD activity 72 hrs post AlkBH8 KD (Figure 6F), supporting the ability of AlkBH8 i-KD to genetically replicate the broad selenoprotein inhibitory mechanism of (S)-RSL3 and auranofin. To further investigate the therapeutic potential of AlkBH8 in CRC the i-KD cell lines were maintained in doxycycline media to assess growth and colony formation. A significant decrease in growth and colony formation was observed across all tested CRC cell lines, with rescue observed by 10mM NAC co-treatment (Figure 6G, S6E-F), supporting our hypothesis that AlkBH8 is a novel therapeutic target in CRC through its ability to induce oxidative stress via modulation of the selenoproteome.

### AlkBH8 is required for CRC xenograft growth and survival *in-vivo*

To further evaluate the effects of AlkBH8 i-KD in an *in-vivo* setting, we chose two of our top responding cell lines (DLD1/SW480) for xenograft implantation. Both shAlkBH8 i-KD cell lines (sh3/sh4) and shNT control cells were implanted (1 x 10^6^/mouse) into the flank of NOD-SCID mice. All mice were fed doxycycline chow (400 mg/kg) beginning on d1, and a subgroup of the mice was additionally co-administered drinking water containing NAC (20mM). No alternative food or water sources were provided while doxycycline chow and NAC water were continually available *ad-libitum* throughout the study. In the DLD1 and SW480 cell lines, the shAlkBH8-3 and shAlkBH8-4 xenografts displayed a decrease in growth and a >50% reduction in final tumor mass as compared to the shNT control. All xenografts displayed a complete rescue when mice were co-administered NAC (Figure 7 A-D). TUNEL (terminal deoxynucleotidyl transferase dUTP nick end labeling) staining was performed to assess endpoint cell death in xenograft tissue, where we observed significantly increased cell death in the shAlkBH8 xenografts as compared to the shNT control. NAC co-treatment fully rescued the increase in cell death, demonstrating that in both cell lines AlkBH8 KD induced oxidative cell death in an *in-vivo* setting (Figure 7E). Next, BrdU (bromodeoxyuridine) staining was performed to assess endpoint cellular proliferation in xenograft tissue. The BrdU positive cell fraction significantly decreased in the shAlkBH8 xenografts as compared to the shNT control, with a full rescue observed upon NAC co-treatment (Figure 7F). These data demonstrate that not only does AlkBH8 KD induce oxidative cell death *in-vivo*, it is also capable of reducing the fraction of proliferating cells across multiple shRNA constructs and two distinct CRC cell lines.

**Figure 7:**
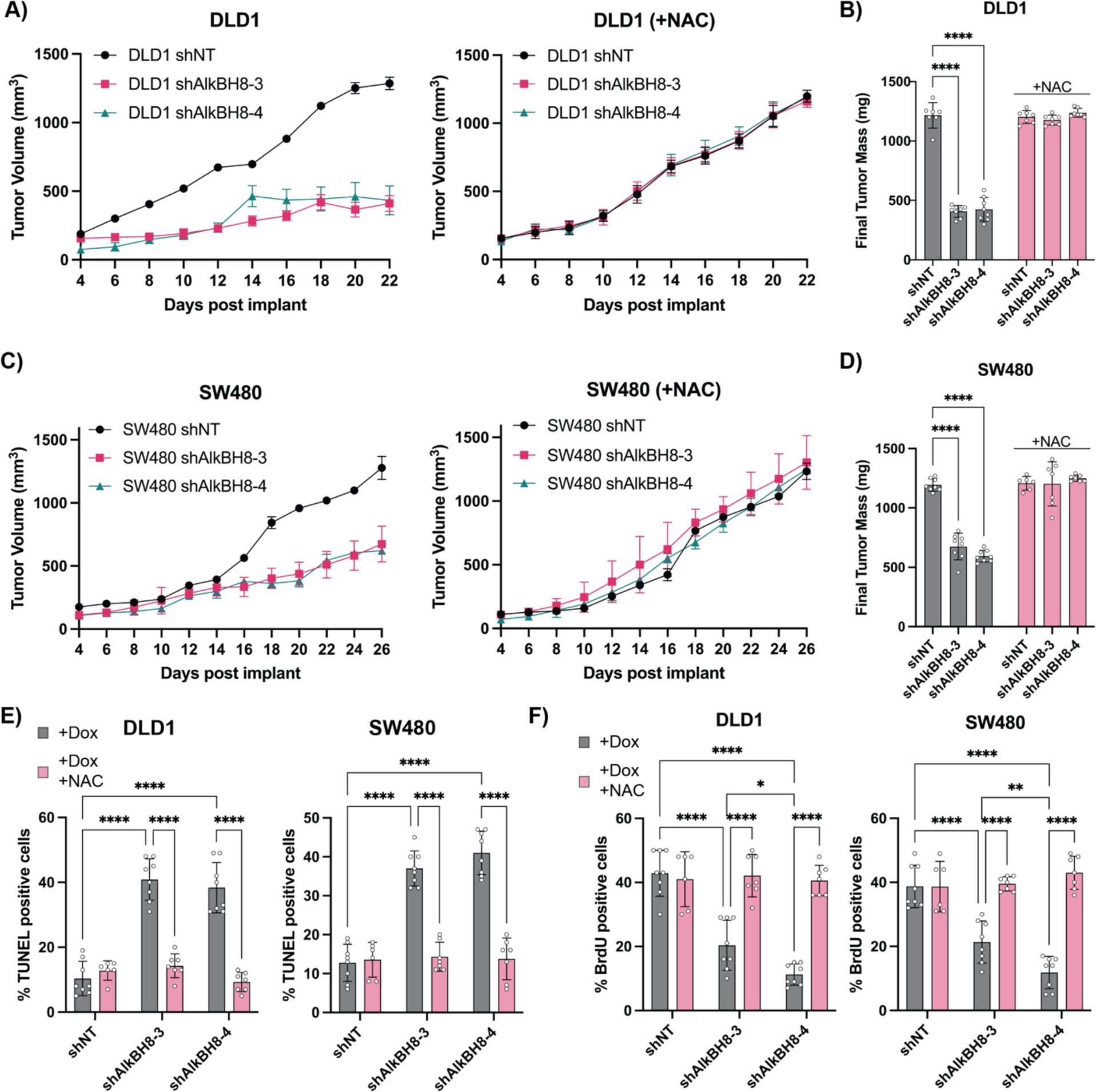
AlkBH8 i-KD induces oxidative stress dependent cell death *in-vivo*. **A)** Tumor volume measurements taken by caliper of DLD1 shNT, shAlkBH8-3, and shAlkBH8-4 xenografts in NOD/SCID mice treated with doxycycline chow +/- co-administration of NAC drinking water. **B)** Final tumor mass of DLD1 xenografts from A. **C)** Tumor volume measurements taken by caliper of SW480 shNT, shAlkBH8-3, and shAlkBH8-4 xenografts in NOD/SCID mice treated with doxycycline chow co-administration of NAC drinking water. **D)** inal tumor mass of SW480 xenografts from C. **E)** Analysis of TUNEL staining (% positive cells) of DLD1 and SW480 xenografts +/- NAC co-treatment. **F)** Analysis of BrdU staining (% positive cells) of DLD1 and SW480 xenografts

## Discussion

Ferroptosis represents a targetable mechanism with high potential for cancer therapeutics. Yet, ferroptosis research has predominantly relied on investigations using small molecules, which can be confounded by unknown off-target effects. In this study we observe that despite broad sensitivity to the GPx4 inhibitor (S)-RSL3, not all CRC cell lines are sensitive to genetic depletion of GPx4, suggesting that at least some of the potency of (S)-RSL3 may stem from GPx4-independent activity. Consequently, our findings shed light on the reduced potency observed with the synthesis of next-generation GPx4 inhibitors such as JKE1674, where heightened target engagement may paradoxically diminish activity. While RSL3 remains a potent ferroptosis inducer, our data adds to its characterization and further cements that it is not solely a GPx4 inhibitor.

While a large fraction of ferroptosis research still focuses on the central role of GPx4 as a lipid peroxide detoxifying enzyme, our study suggests that the broader selenoproteome might equally contribute to redox regulation. In addition to glutathione peroxidases, cells possess a secondary method of peroxide detoxification, the peroxiredoxins (PRDXs) which rely on the selenoprotein TxnRD1 for redox mediated turnover. Our data validate (S)-RSL3 as an inhibitor of TxnRD1 and have led to observations that the DLD1 cell line has ∼2-3-fold higher TxnRD activity than other tested CRC cell lines. Overall, these data lead to a hypothesis that the DLD1 cell line may more heavily utilize the thioredoxin peroxidase system for regulation of intracellular peroxides, perhaps explaining the observed insensitivity of these cells to JKE1674 and GPx4 i-KD.

Recharacterization of the mechanism of (S)-RSL3 from a targeted inhibitor of GPx4 to a broad inhibitor of the selenoproteome has revealed that (S)-RSL3 dually inhibits a binary system of redox regulation in CRC cells, wherein the Txn and GPx systems are both essential to prevent the toxic byproducts of ROS addiction. In line with this hypothesis, we observe that gold therapy using the small molecule auranofin induces oxidative stress in CRC which can lead to ferroptosis, with the exact mechanism of cell death occurring in a highly cell line and dose dependent manner. In our studies, auranofin treatment in established tumors leads to reduced tumor growth with observations of intra-tumoral necrosis suggesting that gold-based inhibition of the selenoproteome can induce tumor cell death *in-vivo*. Furthermore, our studies of low dose auranofin in AOM/DSS models of CRC demonstrate proof of concept that selenoprotein inhibition can limit CRC progression, however due to the on-target toxicities of gold therapy this approach has not yet been tested in the clinic.

These studies have revealed a novel therapeutic target in CRC: the tRNA methyltransferase AlkBH8. Our data demonstrate the essential role of AlkBH8 for the growth of multiple CRC cell lines. Supporting this data, whole-body knockout mice lacking AlkBH8 exhibit no discernible defects, and individuals with homozygous AlkBH8 mutations show a non-lethal phenotype, indicating tolerability to complete loss of AlkBH8 activity^57,66,67^. Importantly, AlkBH8 features two druggable domains, an AlkB Fe(II)/α-ketoglutarate-dependent dioxygenase domain and a methyltransferase domain. Although full-length recombinant AlkBH8 and its substrates remain poorly characterized, we anticipate that their development will facilitate small molecule inhibitor discovery, potentially yielding novel targeted therapies for CRC.

In conclusion, our exploration into the selenoproteome highlights the therapeutic potential of this underexplored protein class. Selenium, and by extension selenocysteine, is orders of magnitude more efficient in reacting with and detoxifying ROS as compared to the sulfur containing cysteine^68^, coinciding with the majority of selenoproteins functioning as antioxidant enzymes in the regulation of oxidative stress. However, only a fraction of the selenoproteome has been investigated to decipher their roles in human biology, and even less for their involvement in cancer development and progression. While our approaches employing AlkBH8 KD, gold therapy, and chloroacetamide warheads to target the selenoproteome yield non-selective inhibition of the selenoproteome, we posit that deeper investigations into the biology of selenoproteins may not only advance our understanding of cellular redox regulation, but also unveil novel therapeutic targets for the treatment of human disease.

## Supporting information

Extended Data 1 - Synthetic Details

## Acknowledgements

The authors would like to acknowledge the members of the Shah and Koutmos laboratories for project feedback and manuscript editing. This work was funded by NIH grants R01CA148828, R01CA245546, and R01DK095201 (Y.M.S.), UMCCC Core grant P30CA046592 (Y.M.S.), CA286898 and GM117141 (M.K.), NIH/NIGMS grant R01GM132129 (J.A.P.), NIH grant R01DK124384 to JDM, and R01CA262439 (Z.T.S.). Sumeet Solanki was supported by a Crohn’s and Colitis Foundation Research fellow award (623914) and the American Heart Association postdoctoral fellowship (19POST34380588). JDM holds patents for methods for modulating ferritinophagy. JDM reports research support from Novartis and Casma Therapeutics and has consulted for Third Rock Ventures and Skyhawk Therapeutics, all unrelated to the work

## Author Contributions

SLD, MK, and YMS initiated the project, performed or directed all experiments, and wrote the manuscript. SD assisted with the majority of experiments as an undergraduate assistant. SS was heavily involved in training and assistance for in-vivo studies. LZ, MOD, NKD, CC, HNB, YZ, NJR, EM, JH, ZS, and IT assisted with in-vitro, bioinformatic, and/or in-vivo experiments. YQ and BWLN assisted with CETSA experiments. MS, JP, and JM performed Mass Spectrometry studies. AB and NN synthesized RSL3 derivatives. MK and YMS provided funding and mentorship for SLD throughout the project.

---

**Figure S1.**
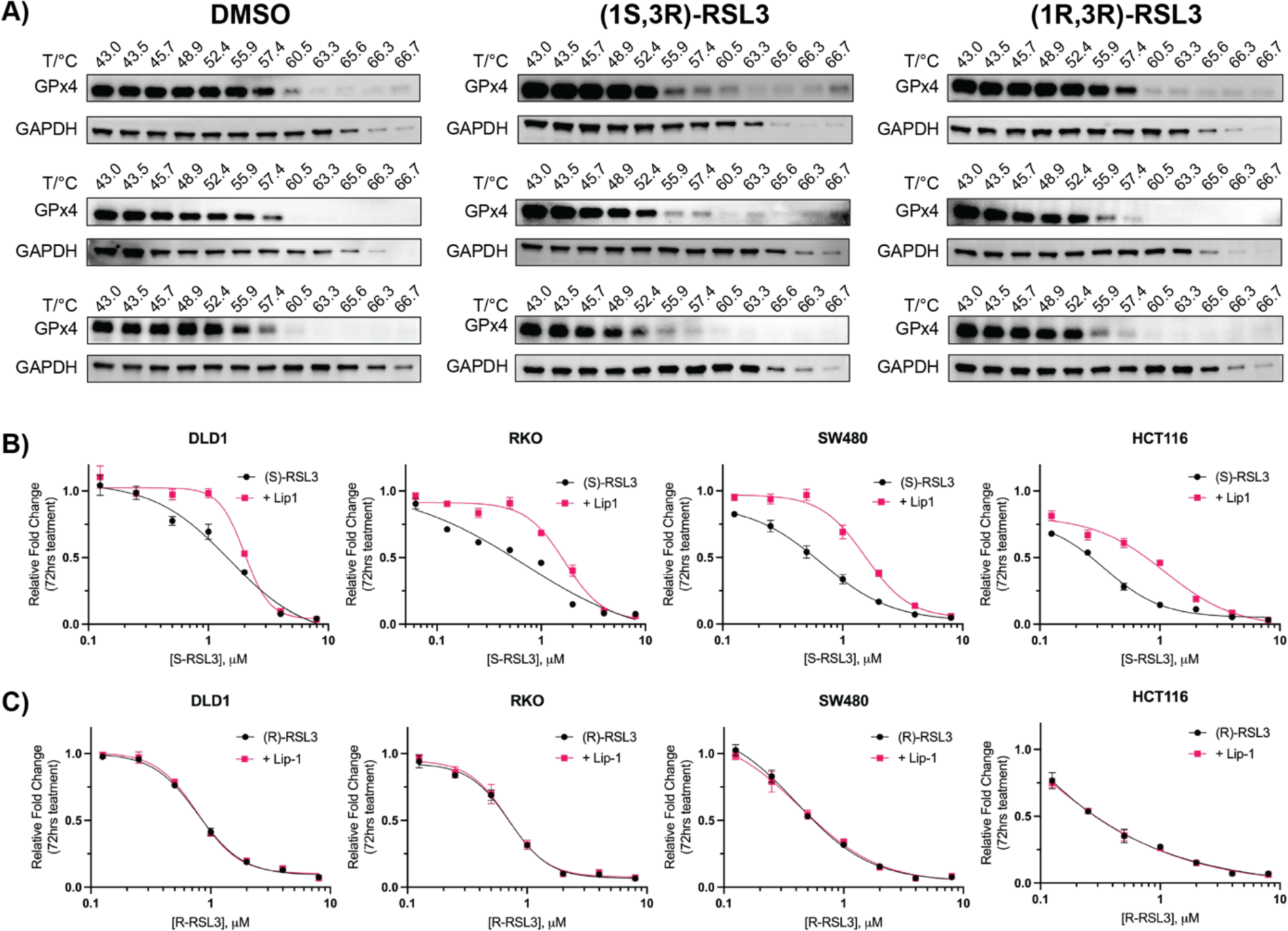
**A)** Western blots of GPx4 CETSA experiments used for generation of Figure 1C. 293T cells were treated with compound as indicated (10μM, 1hr) and lysates were heated to the indicated temperature before pelleting precipitated proteins **B and C)** Cell growth normalized to untreated control at 72 hr following (S) or (R) RSL3 dose response +/- Liproxstatin-1 (1μM) co-treatment in CRC cell lines.

**Figure S2.**
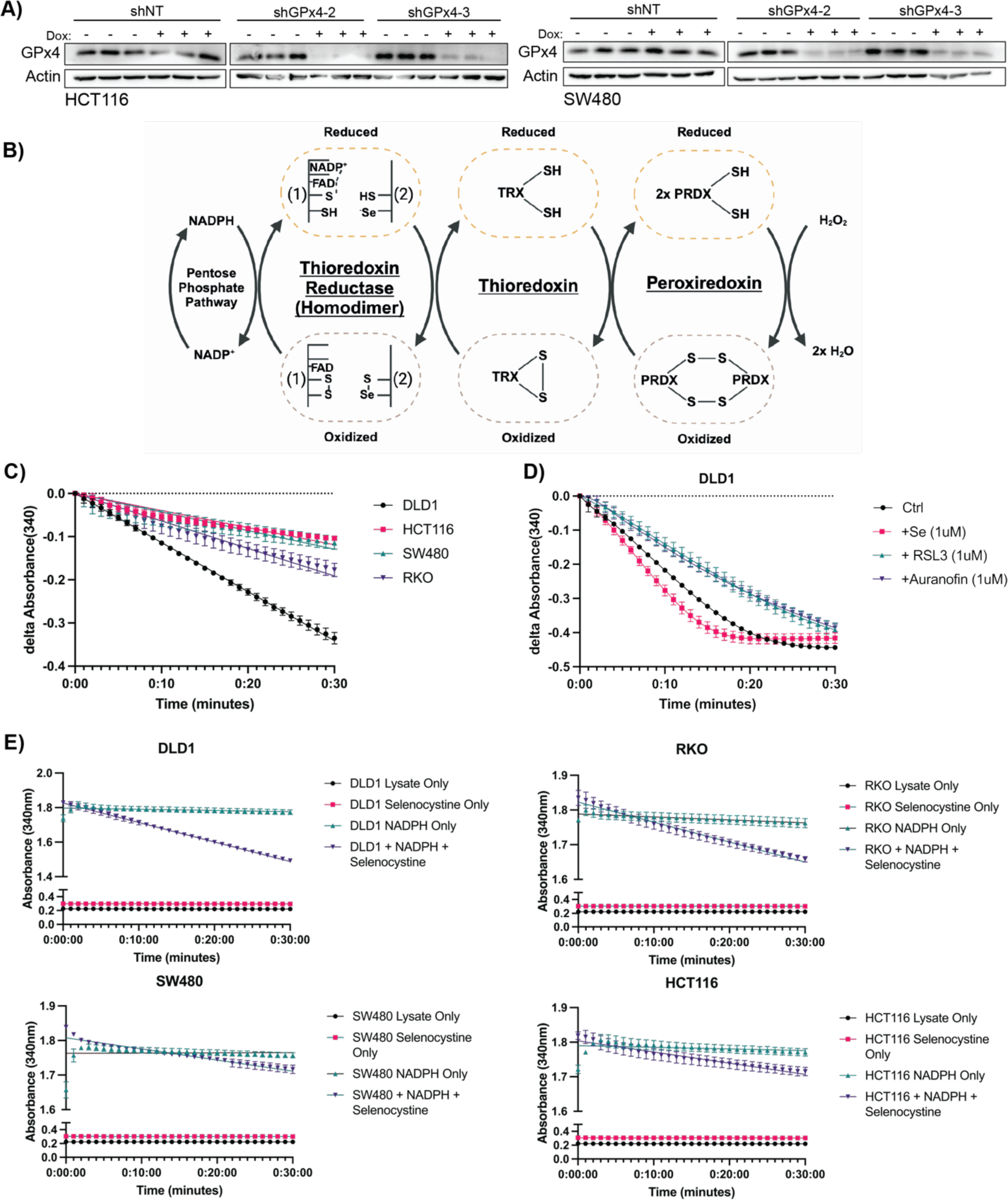
**A)** Western Blots of GPx4 levels in HCT116 and SW480 shNT, shGPx4-2, shGPx4-3 cell lines following 72hr treatment with doxycycline (250 ng/mL) **B)** Schematic of the mechanism of the Thioredoxin Reductase-Peroxiredoxin system, where NADPH from the Pentose Phosphate Pathway is utilized in the selenoprotein Thioredoxin Reductase to recycle oxidized 2-Cys Peroxiredoxins after the redox mediated detoxification of hydrogen peroxide, forming a Peroxiredoxin dimer. The peroxiredoxin dimer is reduced through oxidation of thioredoxin, and oxidized thioredoxin is reduced by TxnRD1 **C)** Thioredoxin Reductase Kinetic Activity Assay measuring NADPH consumption through reduction of NADPH absorbance at 340 nm upon addition of NADPH and Selenocystine to cell lysates of indicated CRC cell lines **D)** Modulation of the thioredoxin reductase activity assay after 24hr treatment of DLD1 cells as indicated. **E)** Internal controls utilized for the thioredoxin reductase kinetic activity assay. Absorbance at 340 nm was measured every 60 s for 30 min on a Cytation 5 plate reader. Wells contained either lysate only, lysate + selenocystine, lysate + NADPH, or lysate + selenocystine & NADPH as indicated. 3 technical replicates were averaged for each point and plotted as mean +/- SD

**Figure S3.**
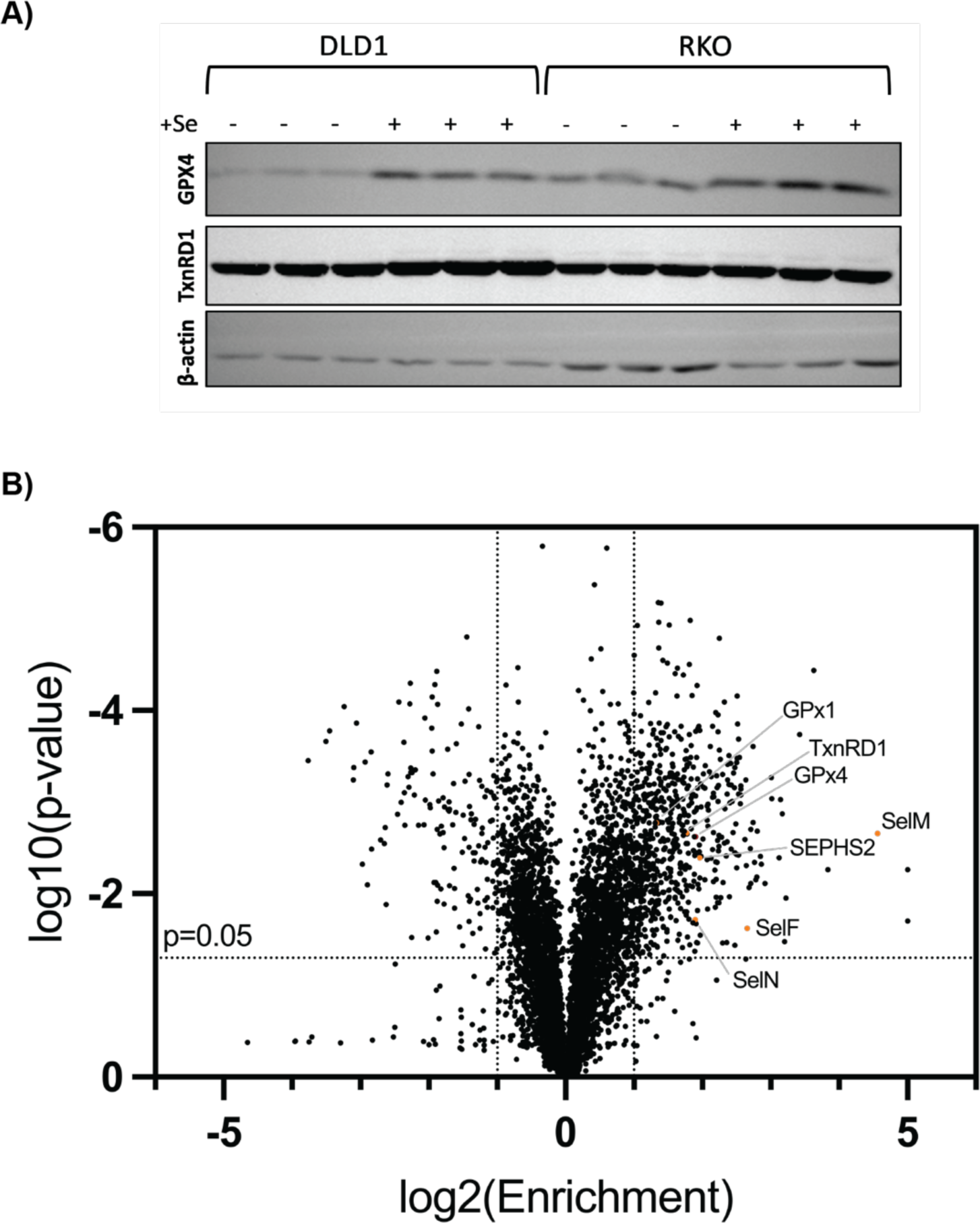
**A)** Western blot analysis of GPx4 and TxnRD1 protein levels in DLD1 and RKO cell lines following 24 hr supplemention with 1 μM Sodium Selenite. **B)** AP-MS results of Biotin-Cpd24 pulldown compared to Cpd24 mock pulldown control. Significant selenoproteins hits are identified

**Figure S4.**
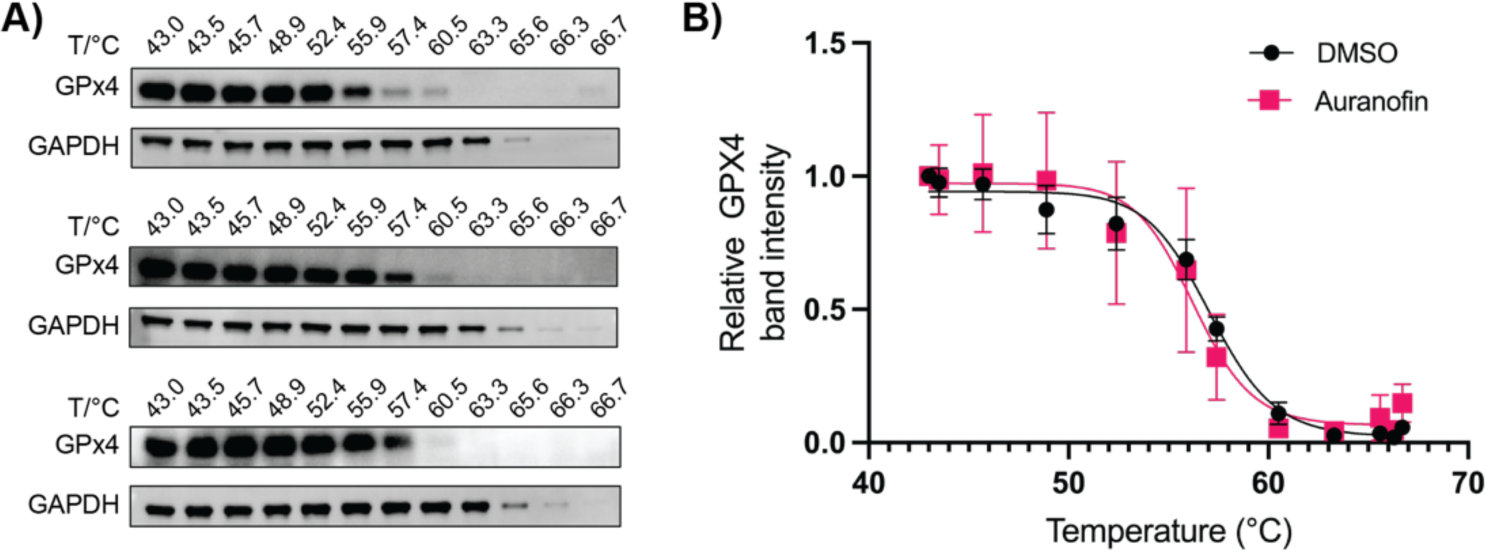
**A)** CETSA analysis of GPx4 protein thermal stability following auranofin treatment (10 μM, 1 hr) in 293T cells. DMSO controls are included as Figure S1A. **B)** Results of CETSA analysis from A calculating deviation of GPx4 band intensity between DMSO treated and auranofin treated cells (DMSO controls shown as Fig S1A)

**Figure S5.**
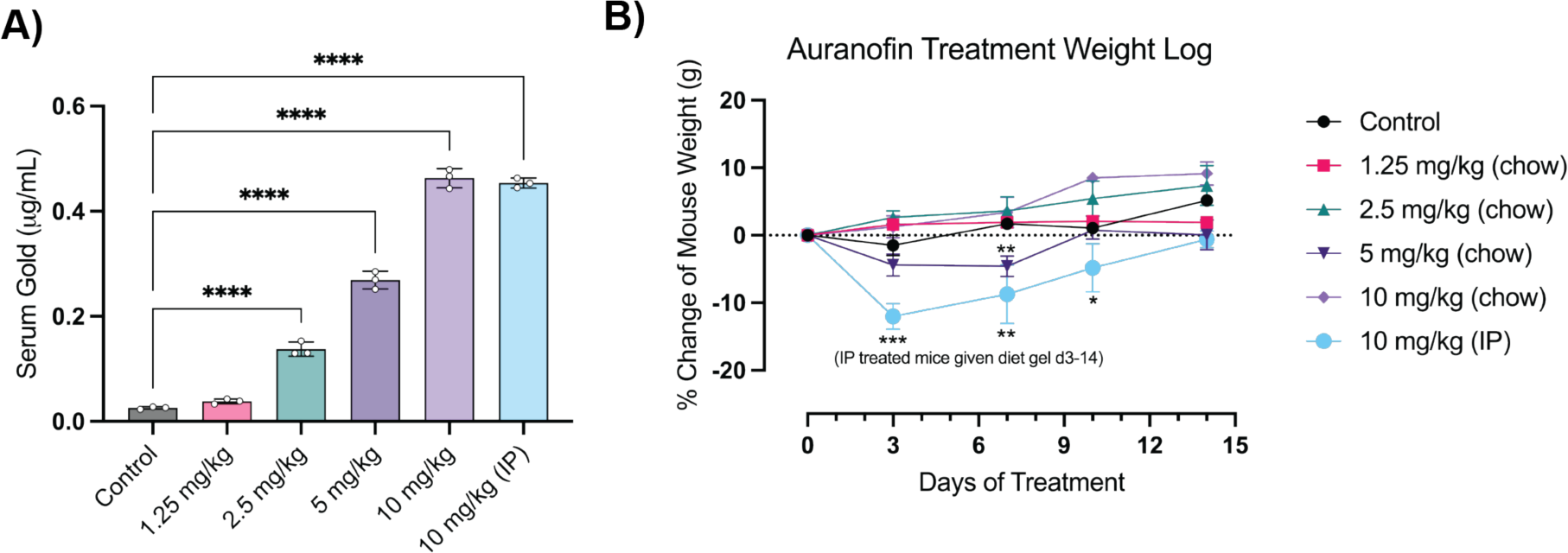
**A)** Terminal serum gold measurements by ICPMS of non tumor bearing control mice (C57Bl/6J) treated with indicated dosing of auranofin via chow or IP as indicated for two weeks. **B)** Weight log of non tumor bearing control mice treated with auranofin via chow or IP as indicated. IP treated mice were administered supplemental diet gel beginning d3 following rapid initial weight loss

**Figure S6.**
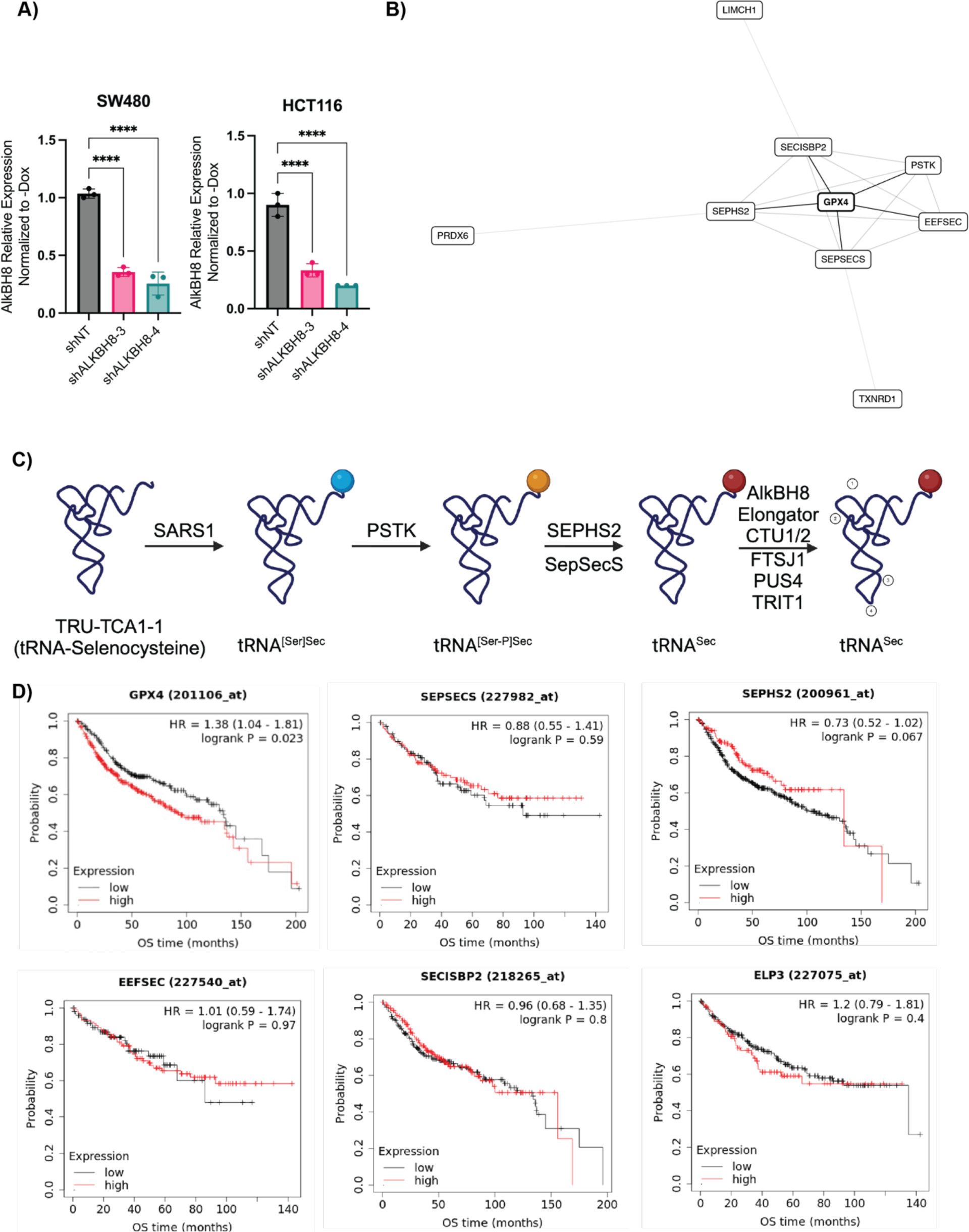
**A)** qPCR analysis of shNT or shAlkBH8 HCT116/SW480 cell lines measuring AlkBH8 mRNA expression following 72 hr dox treatment normalized to -dox control. **B)** CRISPR co-essentiality network map of GPx4 co-dependencies with distances indicating degree of co-essentiality. **C)** Biosynthetic pathway of tRNA-Selenocysteine where following transcription of the TRU-TCA1-1 gene, the tRNA-Sec mRNA undergoes on-tRNA biosynthesis of selenocysteine and post-transcriptional modification. **D)** Km plots of select genes in the tRNA-sec pathway with sufficient patient samples to perform a high-powered analysis

