## Extended Data 1 - Synthetic Details for "Recharacterization of RSL3 reveals that the selenoproteome is a druggable target in colorectal cancer"

**Scheme 1. Synthesis of RSL3 and Biotinylated Analog^a^**

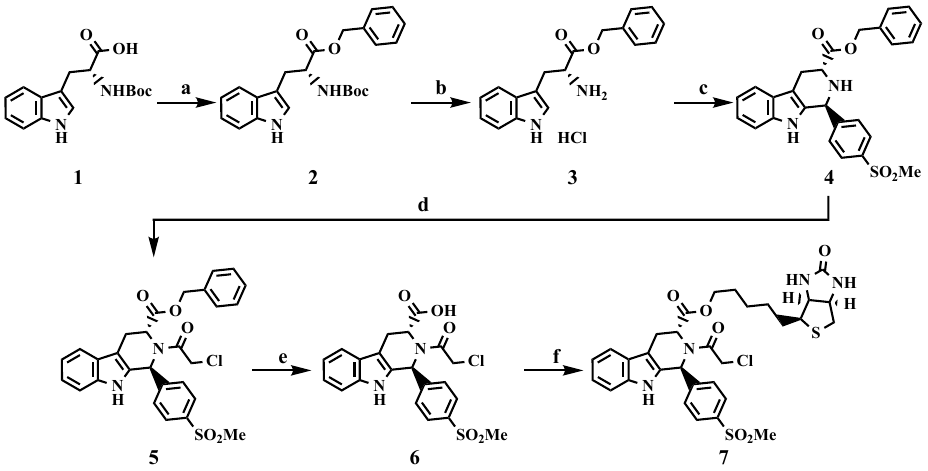

^a^Reagents and conditions: (a) BnOH, EDC, DMAP, THF, rt, overnight; (b) 4 N HCl in 1,4-dioxane, rt, overnight, 92% over 2 steps; (c) 4-(methylsulfonyl)benzaldehyde, IPA, reflux, 87%; (d) ClCH_2_C(O)Cl, TEA, CH_3_CN, reflux, 87%; (e) H_2_, Pd/C, EtOH, rt, overnight, 72%; (f) *D*-biotinol, EDC, DMAP, THF, rt, overnight, 4%.

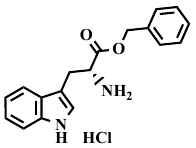
(AGB-312a) *D*-Benzyl tryptophanate hydrochloride 3.

To a solution of (*tert*-butoxycarbonyl)-*D*-tryptophan (1.215 g, 4 mmol) in anhydrous THF (9 mL) were added DMAP (975.4 mg, 8 mmol), benzyl alcohol (0.83 mL, 8 mmol) and EDC (3.06 g, 16 mmol). The reaction mixture was then stirred at rt for overnight. The solvent was removed under reduced pressure. The residue was dissolved in СH_2_Cl_2_ and washed with NaOH (1 N), HCl (1 N) dried over anhydrous MgSO_4_, filtered and concentrated under reduced pressure. The residue was treated with 4 N HCl in 1,4-dioxane (6 mL) and stirred at rt overnight. The reaction mixture was concentrated under reduced pressure to dryness and purified by suspending in anhydrous acetone, filtered, washed with acetone and dried to give **3** (1.2159 g, 92% over 2 steps) as a white solid. ^1^H NMR (599 MHz, DMSO-*d_6_*) δ 11.09 – 11.08 (m, 1H), 8.57 (brs, 3H), 7.51 (dd, *J* = 7.9, 1.0 Hz, 1H), 7.39 (dt, *J* = 8.2, 1.0 Hz, 1H), 7.35 – 7.31 (m, 3H), 7.22 – 7.18 (m, 3H), 7.10 (ddd, *J* = 8.2, 7.0, 1.2 Hz, 1H), 7.00 (ddd, *J* = 8.0, 7.0, 1.1 Hz, 1H), 5.14 (d, *J* = 12.4 Hz, 1H), 5.06 (d, *J* = 12.4 Hz, 1H), 4.31 – 4.29 (m, 1H), 3.35 – 3.32 (m, 1H), 3.28 (dd, *J* = 15.0, 6.9 Hz, 1H).

**(AGB-313-2) Benzyl (1*S*,3*R*)-1-(4-(methylsulfonyl)phenyl)-2,3,4,9-tetrahydro-1*H*-pyrido[3,4-*b*]indole-3-carboxylate 4.**

A mixture of *D*-benzyl tryptophanate hydrochloride (616.5 mg, 1.86 mmol) and 4-(methylsulfonyl)benzaldehyde (411.9 mg, 2.2 mmol) in isopropyl alcohol (12 mL) was heated and stirred at 80 °C. Cooled to ambient and collected the solid product to give **4** (742.9 mg, 87%) as a white solid. The filtrate was evaporated under reduced pressure. The residue was dissolved in СH_2_Cl_2_ and washed with NaHCO_3_ dried over anhydrous MgSO_4_, filtered and concentrated. The residue was purified by column chromatography on silica gel using MeOH/CH_2_Cl_2_/NH_3_ 1/50/0.1 as eluent to give the title compound as a white solid.
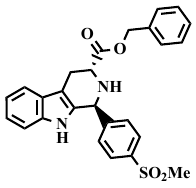
^1^H NMR (599 MHz, CDCl_3_) δ 7.90 – 7.87 (m, 2H), 7.59 – 7.54 (m, 2H), 7.52 – 7.49 (m, 2H), 7.31 – 7.24 (m, 3H), 7.19 (td, *J* = 7.6, 1.4 Hz, 1H), 7.15 (ddd, *J* = 8.2, 7.2, 1.2 Hz, 1H), 5.50 (t, *J* = 1.7 Hz, 1H), 5.16 (q, *J* = 12.4 Hz, 2H), 3.98 (t, *J* = 5.8 Hz, 1H), 3.29 (ddd, *J* = 15.4, 5.5, 1.6 Hz, 1H), 3.21 (ddd, *J* = 15.4, 6.2, 1.6 Hz, 1H), 3.04 (s, 3H).

(AGB-313a2) Benzyl (1S,3R)-2-(2-chloroacetyl)-1-(4-(methylsulfonyl)phenyl)-2,3,4,9-tetrahydro-1*H*-pyrido[3,4-*b*]indole-3-carboxylate 5.

To the mixture of **4** (742.9 mg, 1.6 mmol) with triethylamine (0.67 mL, 4.84 mmol) in anhydrous acetonitrile (4 mL), chloroacetyl chloride (0.2 mL, 2.42 mmol) was carefully added at rt. The resulting mixture was heated for 1 min, cooled to rt, evaporated under reduced pressure and purified by column chromatography on silica gel using MeOH/CH_2_Cl_2_ 1/100 as eluent to give the title compound **5** (742.9 mg, 95%) as a white solid.
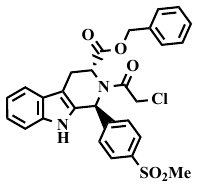
^1^H NMR (599 MHz, CDCl_3_) δ 8.30 (brs, 1H), 7.73 (d, *J* = 8.2 Hz, 2H), 7.49 (d, *J* = 7.4 Hz, 1H), 7.43 (d, *J* = 8.2 Hz, 2H), 7.22 – 7.06 (m, 8H), 6.94 (d, *J* = 7.7 Hz, 2H), 6.09 (s, 1H), 5.26 (d, *J* = 4.9 Hz, 1H), 5.13 – 5.00 (m, 3H), 4.15 (d, *J* = 13.2 Hz, 1H), 4.05 – 4.00 (m, 1H), 3.79 (d, *J* = 15.1 Hz, 1H), 3.50 (dd, *J* = 15.3, 5.2 Hz, 1H), 2.93 (s, 3H).

**(AGB-314) (1*S*,3*R*)-2-(2-Chloroacetyl)-1-(4-(methylsulfonyl)phenyl)-2,3,4,9-tetrahydro-1*H*-pyrido[3,4-*b*]indole-3-carboxylic acid 6.**

The mixture of **5** (213.4 mg, 0.4 mmol) with Pd/C (162 mg) in ethanol (12 mL) was stirred under hydrogen atmosphere overnight at room temperature. Suspension was filtered through Celite evaporated under reduced pressure. The residue was purified by column chromatography on silica gel using MeOH/CH_2_Cl_2_ 1/10 as eluent to give the title compound 6a (127.8 mg, 72%) as a white solid.
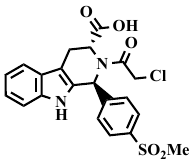
^1^H NMR (599 MHz, CD_3_OD) δ 7.79 (d, *J* = 8.4 Hz, 2H), 7.65 (d, *J* = 8.4 Hz, 2H), 7.47 (d, *J* = 7.7 Hz, 1H), 7.19 (d, *J* = 8.0 Hz, 1H), 7.03 (t, *J* = 7.4 Hz, 1H), 6.99 (t, *J* = 7.4 Hz, 1H), 6.19 (brs, 1H), 5.25 (brs, 1H), 4.53 (d, *J* = 13.7 Hz, 1H), 4.21 (d, *J* = 13.7 Hz, 1H), 3.75 (d, *J* = 15.0 Hz, 1H), 3.47 – 3.41 (m, 1H), 2.97 (s, 3H).

LCMS: r.t. 4.327, 100%; 446.9, 448.9 [M+H].

(AGB-315a) 5-((3a*S*,4*S*,6a*R*)-2-Oxohexahydro-1*H*-thieno[3,4-*d*]imidazol-4-yl)pentyl (1*S*,3*R*)-2-(2-chloroacetyl)-1-(4-(methylsulfonyl)phenyl)-2,3,4,9-tetrahydro-1*H*-pyrido[3,4-*b*]indole-3-carboxylate 7.

To a solution of 6 (74.9 mg, 0.1676 mmol) in anhydrous THF (4 mL) were added DMAP (40.9 mg, 0.335 mmol), *D*-biotinol (38.6 mL, 0.1676 mmol) and EDC (128.5 mg, 0.67 mmol). The reaction mixture was then stirred at rt for overnight. The solvent was removed under reduced pressure. The residue was dissolved in СH_2_Cl_2_ and washed with HCl (1 N) dried over anhydrous MgSO_4_, filtered and concentrated under reduced pressure. The residue was purified by preparative plate chromatography on silica gel using MeOH/CH_2_Cl_2_ 1/25 as eluent to give the title compound 7 (4.5 mg, 4%) as a colorless oil.
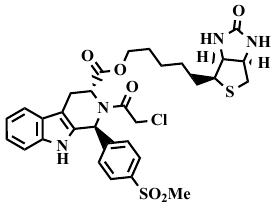
^1^H NMR (599 MHz, CDCl_3_) δ 9.92 (brs, 1H), 9.42 (brs, 1H), 7.91 (d, *J* = 8.2 Hz, 2H), 7.82 (d, *J* = 8.2 Hz, 2H), 7.79 (d, *J* = 7.8 Hz, 2H), 7.66 (d, *J* = 8.2 Hz, 2H), 7.46 (d, *J* = 8.0 Hz, 1H), 7.40 (d, *J* = 7.9 Hz, 1H), 7.32 (d, *J* = 8.2 Hz, 2H), 7.11 (ddd, *J* = 8.3, 6.9, 1.3 Hz, 2H), 7.05 (q, *J* = 7.7 Hz, 2H), 6.94 (brs, 1H), 5.42 (brs, 1H), 5.31 (brs, 1H), 5.26 (brs, 1H), 5.08 (brs, 1H), 4.96 (brs, 1H), 4.91 (brs, 1H), 4.82 (brs, 1H), 4.73 (t, *J* = 11.2 Hz, 1H).

LCMS: r.t. 7.002, 100%, 659.0; 461.0 [M+H]; 657.0; 658.9 [M-H].

**Scheme 2. Synthesis of RSL3 and Biotinylated Analog^a^**

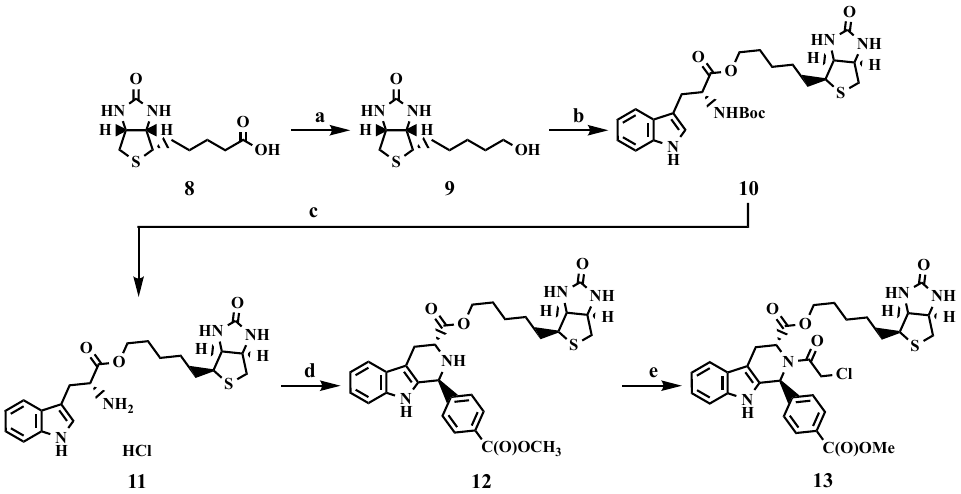

^a^Reagents and conditions: (a) LiAlH_4_, THF, reflux, overnight, 45%; (b) (*tert*-butoxycarbonyl)-*D*-tryptophan, EDC, DMAP, THF, rt, overnight, 74%; (c) 4 N HCl in 1,4-dioxane, rt, overnight, 91%; (d) methyl 4-formylbenzoate, IPA, reflux, 15%; (e) ClCH_2_C(O)Cl, TEA, CH_3_CN, reflux, 19%.

**(AGB-329a) *D*-Biotinol 9.**

To a suspension of *D*-biotin (3.5356 g, 14.47 mmol) in anhydrous THF (60 mL) was added LiAlH_4_ (3.3 g, 86.83 mmol). The reaction mixture was then stirred at 90 °C for overnight. The resulting suspension was cooling to rt and was added water (10 mL), dried over anhydrous MgSO_4_, filtrated, evaporated. The residue was purified by column chromatography on silica gel using MeOH/CH_2_Cl_2_ 1/10 as eluent to provide the title compound **9** (1.5 g, 45%) as a white solid.
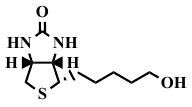
 ^1^H NMR (599 MHz, DMSO-*d_6_*) δ 6.42 (brs, 1H), 6.35 (brs, 1H), 4.43 (t, *J* = 5.1 Hz, 1H), 4.33 – 4.29 (m, 1H), 4.13 (ddd, *J* = 7.8, 4.5, 2.0 Hz, 1H), 3.37 (td, *J* = 6.5, 5.1 Hz, 3H), 3.12 – 3.07 (m, 1H), 2.81 (dd, *J* = 12.5, 5.1 Hz, 1H), 2.57 (d, *J* = 12.4 Hz, 1H), 1.65 – 1.57 (m, 1H), 1.50 – 1.23 (m, 8H).

(AGB-330a) 5-((3a*S*,4*S*,6a*R*)-2-Oxohexahydro-1*H*-thieno[3,4-*d*]imidazol-4-yl)pentyl (*tert*-butoxycarbonyl)-*D*-tryptophanate 10.

To a solution of (*tert*-butoxycarbonyl)-*D*-tryptophan (408.2 mg, 1.34 mmol) in anhydrous DMF (6 mL) were added DMAP (327.8 mg, 2.68 mmol), *D*-biotinol (309 mg, 1.34 mmol) and EDC (1.0287 g, 5.4 mmol). The reaction mixture was then stirred at rt for overnight. The mixture was then evaporated under reduced pressure, diluted with water and CH_2_Cl_2_, washed with water (3 x 200 mL), dried over anhydrous MgSO_4_, filtered, and concentrated. The residue was purified by column chromatography on silica gel using MeOH/CH_2_Cl_2_ 1/10 as eluent to provide the title compound **10** (492.1 mg, 71%) as a white foam.
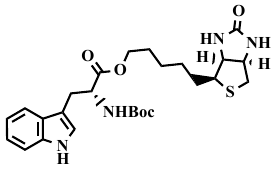
^1^H NMR (599 MHz, CDCl_3_) δ 9.19 (brs, 1H), 7.55 (d, *J* = 7.9 Hz, 1H), 7.38 (d, *J* = 8.1 Hz, 1H), 7.16 (t, *J* = 7.6 Hz, 1H), 7.09 (t, *J* = 7.5 Hz, 1H), 7.02 (brs, 1H), 6.33 (brs, 1H), 5.28 (brs, 1H), 5.16 (d, *J* = 8.3 Hz, 1H), 4.67 – 4.59 (m, 1H), 4.44 (dd, *J* = 7.9, 4.9 Hz, 1H), 4.25 (dd, *J* = 8.3, 4.7 Hz, 1H), 4.03 – 3.89 (m, 2H), 3.34 – 3.20 (m, 2H), 3.07 (td, *J* = 7.5, 4.6 Hz, 1H), 2.89 – 2.84 (m, 1H), 2.67 (d, *J* = 12.8 Hz, 1H), 1.77 (brs, 1H), 1.56 (q, *J* = 7.8 Hz, 2H), 1.44 (s, 9H), 1.38 (d, *J* = 13.3 Hz, 2H), 1.26 (q, *J* = 6.7 Hz, 2H), 1.01 (h, *J* = 8.5 Hz, 1H); ^13^C NMR (151 MHz, CDCl_3_) δ 172.74, 164.36, 155.38, 136.39, 127.87, 123.25, 121.97, 119.40, 118.77, 111.56, 109.80, 79.93, 65.41, 62.34, 60.19, 55.90, 54.71, 40.57, 28.78, 28.59, 28.47, 28.40, 28.11, 25.67.

(AGB-333) 5-((3a*S*,4*S*,6a*R*)-2-Oxohexahydro-1*H*-thieno[3,4-*d*]imidazol-4-yl)pentyl *D*-tryptophanate hydrochloride 11.

The compound **10** (515.7 mg, 0.998 mmol) was treated with 4 N HCl in 1,4-dioxane (6 mL) and stirred at rt overnigth. The reaction mixture was concentrated under reduced pressure to dryness and purified by suspending in anhydrous acetone, filtered, washed with acetone and dried to give **11** (410 mg, 91%) as a white solid.
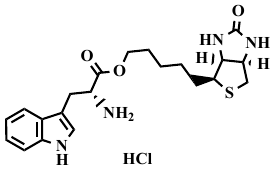
^1^H NMR (599 MHz, DMSO-*d_6_*) δ 11.10 – 11.00 (m, 1H), 8.41 (brs, 3H), 7.50 (d, *J* = 7.9 Hz, 1H), 7.38 (d, *J* = 8.1 Hz, 1H), 7.23 (d, *J* = 2.6 Hz, 1H), 7.10 (t, *J* = 7.6 Hz, 1H), 7.02 (t, *J* = 7.5 Hz, 1H), 6.42 (s, 1H), 6.38 (s, 1H), 4.31 (dd, *J* = 7.7, 5.1 Hz, 1H), 4.23 (t, *J* = 6.6 Hz, 1H), 4.15 – 4.10 (m, 1H), 4.07 – 3.95 (m, 2H), 3.32 – 3.19 (m, 2H), 3.11 – 3.04 (m, 1H), 2.81 (dd, *J* = 12.4, 5.1 Hz, 1H), 2.58 (d, *J* = 12.4 Hz, 1H), 1.56 (ddd, *J* = 11.9, 10.1, 6.1 Hz, 1H), 1.48 – 1.36 (m, 3H), 1.32 – 1.11 (m, 4H).

**(AGB-353a1i)** **5-((3a*S*,4*S*,6a*R*)-2-Oxohexahydro-1*H*-thieno[3,4-*d*]imidazol-4-yl)pentyl (1*S*,3*R*)-1-(4-(methoxycarbonyl)phenyl)-2,3,4,9-tetrahydro-1*H*-pyrido[3,4-*b*]indole-3-carboxylate 12.**

A mixture of **11** (262.9 mg, 0.58 mmol) and methyl 4-formylbenzoate (114.3 mg, 0.7 mmol) in isopropyl alcohol (20 mL) was heated and stirred at 80 °C for 6 h. Cooled mixture was evaporated under reduced pressure. The residue was dissolved in СH_2_Cl_2_ and washed with NaHCO_3_ dried over anhydrous MgSO_4_, filtered and concentrated. The residue was purified by plate chromatography on silica gel using MeOH/CH_2_Cl_2_/NH_3_ 1/50/0.1 as eluent to give the title compound **12** (50.4 mg, 15%) as a white solid
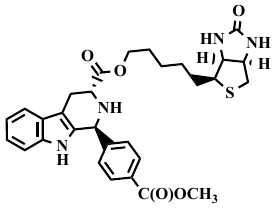
. ^1^H NMR (599 MHz, CDCl_3_) δ 8.03 (d, *J* = 8.2 Hz, 2H), 7.55 (d, *J* = 8.8 Hz, 1H), 7.48 (d, *J* = 8.2 Hz, 2H), 7.46 (s, 1H), 7.21 (d, *J* = 7.1 Hz, 1H), 7.17 – 7.08 (m, 2H), 5.31 (s, 1H), 5.29 (s, 1H), 4.94 (s, 1H), 4.50 – 4.46 (m, 1H), 4.31 – 4.27 (m, 1H), 4.22 (qt, *J* = 10.8, 6.6 Hz, 2H), 3.97 (d, *J* = 4.3 Hz, 1H), 3.95 (d, *J* = 4.3 Hz, 1H), 3.92 (s, 3H), 3.27 – 3.21 (m, 1H), 3.18 – 3.12 (m, 1H), 3.01 (ddd, *J* = 15.1, 11.1, 2.6 Hz, 1H), 2.90 (dd, *J* = 12.9, 5.1 Hz, 1H), 2.70 (d, *J* = 12.7 Hz, 1H), 1.76 – 1.65 (m, 4H), 1.51 – 1.40 (m, 4H).

**(AGB-366I.1) 5-((3a*S*,4*S*,6a*R*)-2-Oxohexahydro-1*H*-thieno[3,4-*d*]imidazol-4-yl)pentyl (1*S*,3*R*)-2-(2-chloroacetyl)-1-(4-(methoxycarbonyl)phenyl)-2,3,4,9-tetrahydro-1*H*-pyrido[3,4-*b*]indole-3-carboxylate 13.**

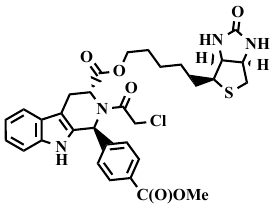
To the mixture of **12** (50.4 mg, 0.09 mmol) with triethylamine (27.2 μL, 0.2687 mmol) in anhydrous acetonitrile (1.5 mL), chloroacetyl chloride (15.2 mg, 0.13 mmol) was carefully added at rt. The resulting mixture was heated for 1 min, cooled to rt, evaporated under reduced pressure and purified by plate on silica gel using EtOAc/Hexanes 1/1 as eluent to give the title compound **13** (10.9 mg, 19%) as a white foam. ^1^H NMR (599 MHz, CDCl_3_) δ 8.30 (brs, 1H), 7.88 (d, *J* = 8.6 Hz, 2H), 7.59 (d, *J* = 7.7 Hz, 1H), 7.36 (d, *J* = 8.2 Hz, 2H), 7.31 (d, *J* = 8.0 Hz, 1H), 7.20 (t, *J* = 7.6 Hz, 1H), 7.16 (t, *J* = 7.5 Hz, 1H), 6.93 (brs, 1H), 5.82 (brs, 1H), 5.26 (brs, 1H), 4.95 (d, *J* = 6.3 Hz, 1H), 4.50 – 4.45 (m, 1H), 4.35 (d, *J* = 12.1 Hz, 1H), 4.27 – 4.20 (m, 2H), 3.88 (s, 3H), 3.78 – 3.66 (m, 2H), 3.19 (ddd, *J* = 15.7, 6.8, 1.8 Hz, 1H), 3.09 (d, *J* = 5.0 Hz, 1H), 3.01 (td, *J* = 7.4, 4.6 Hz, 1H), 2.89 (dd, *J* = 12.9, 5.1 Hz, 1H), 2.69 (d, *J* = 12.7 Hz, 1H), 1.41 (q, *J* = 7.7 Hz, 2H), 1.23 – 1.12 (m, 2H), 1.08 (brs, 23H), 0.92 (brs, 2H).

LCMS: r.t. 7.716, 100%, 639.1; 641.1 [M+H].

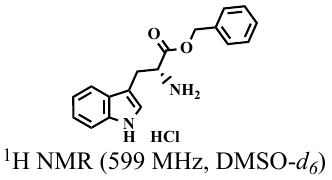

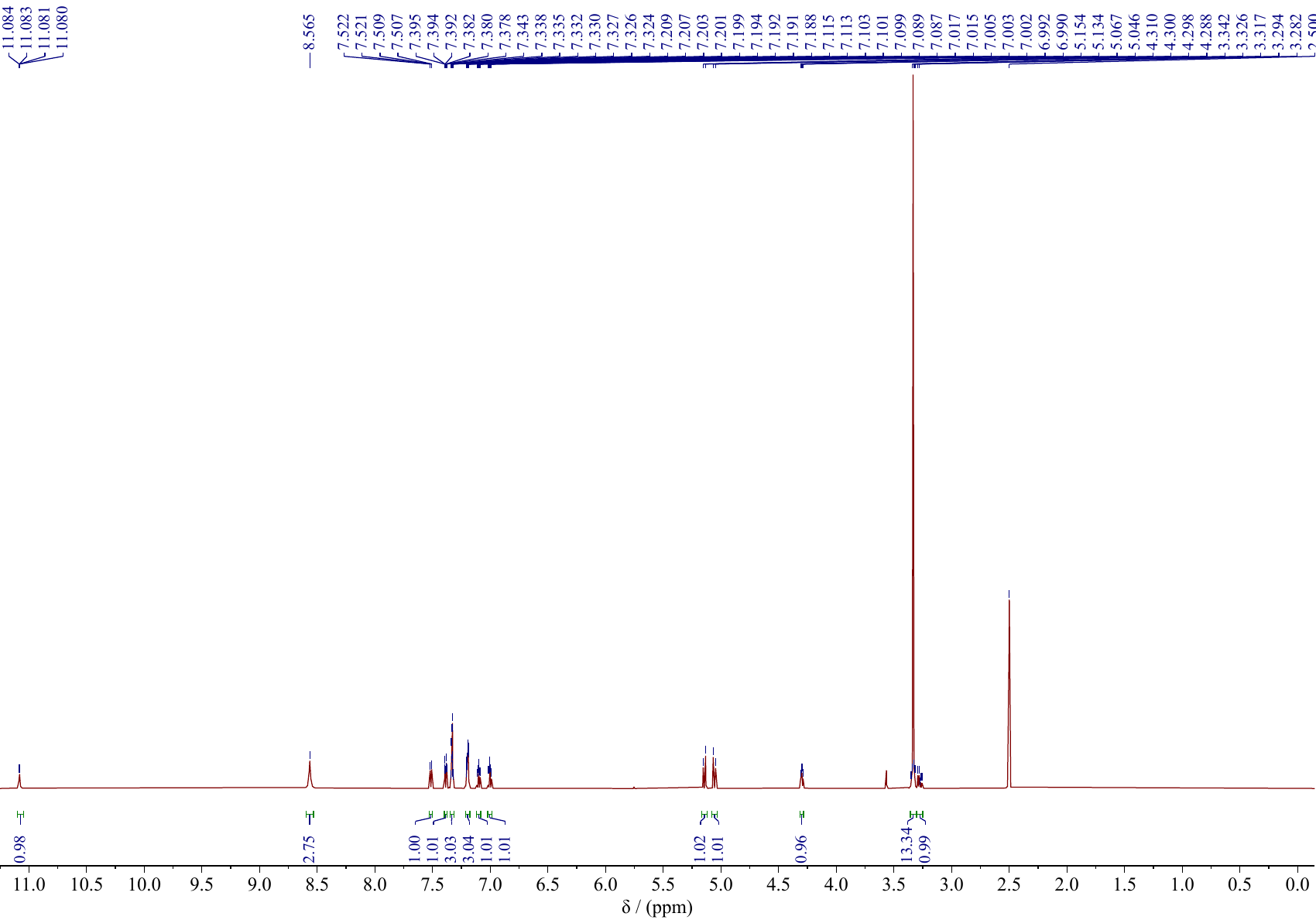

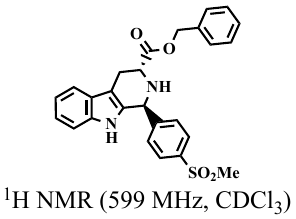

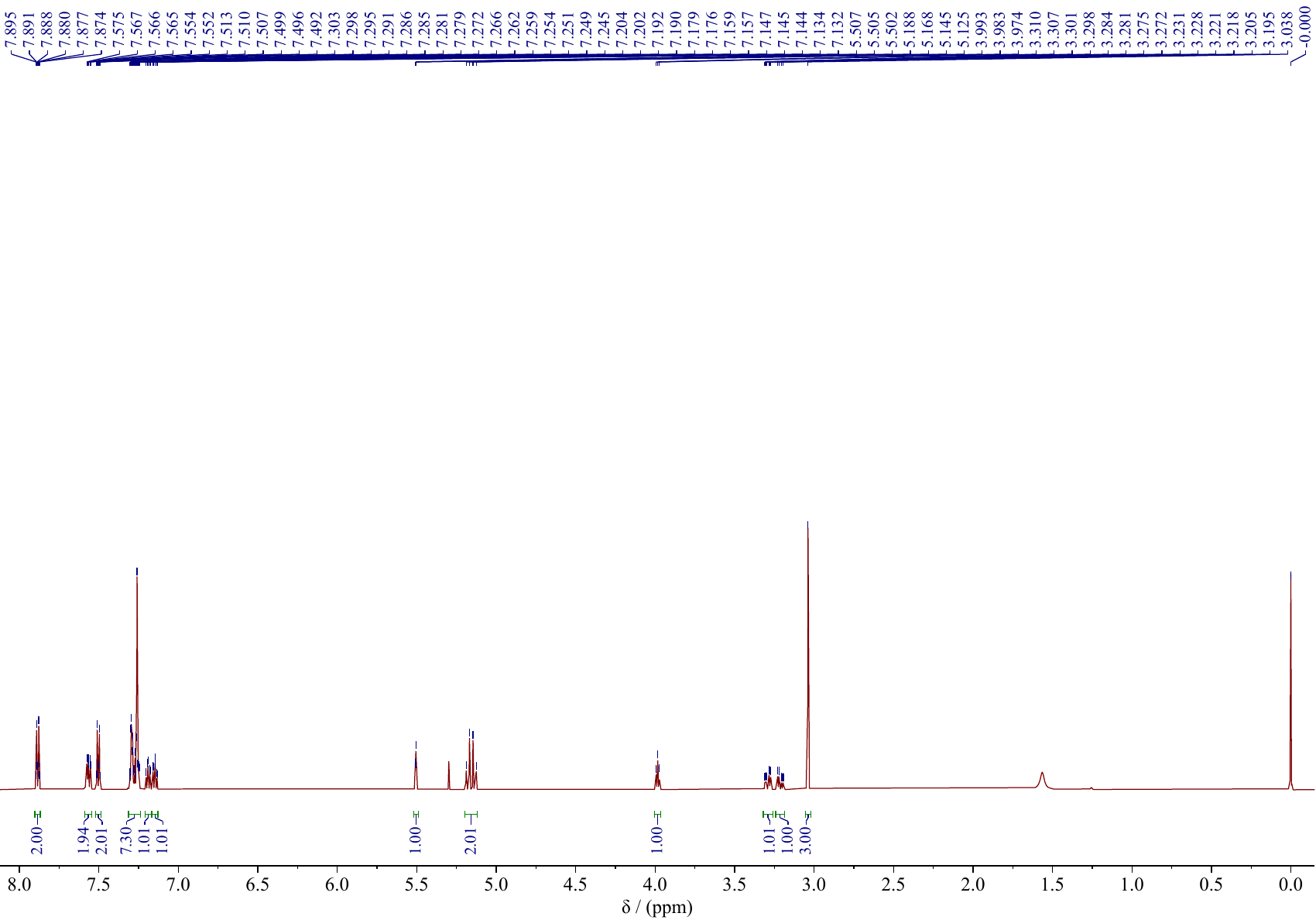

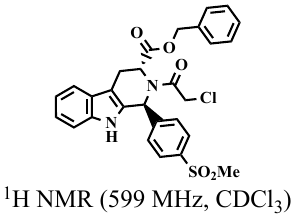

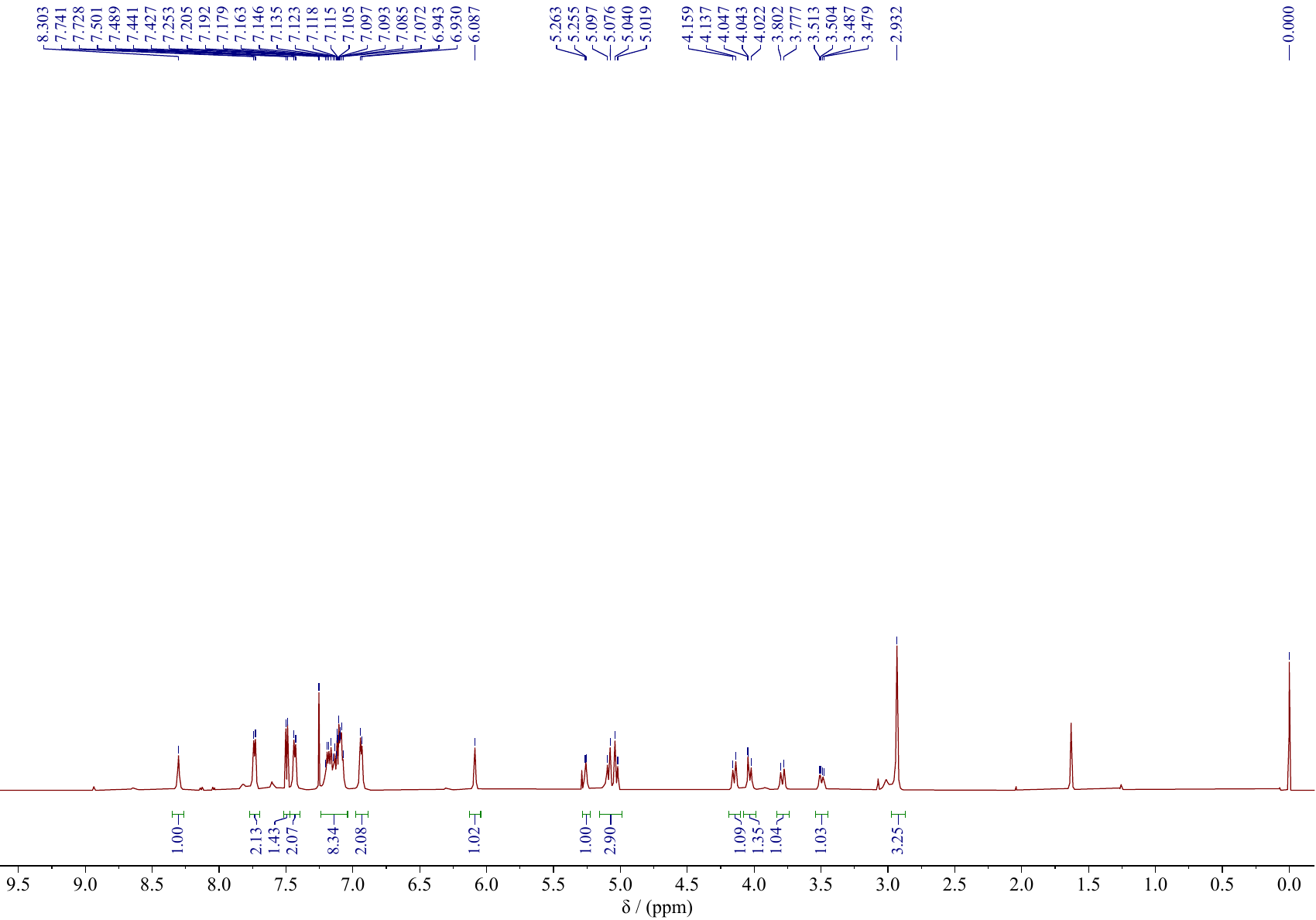

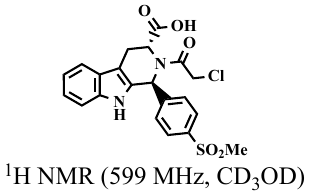

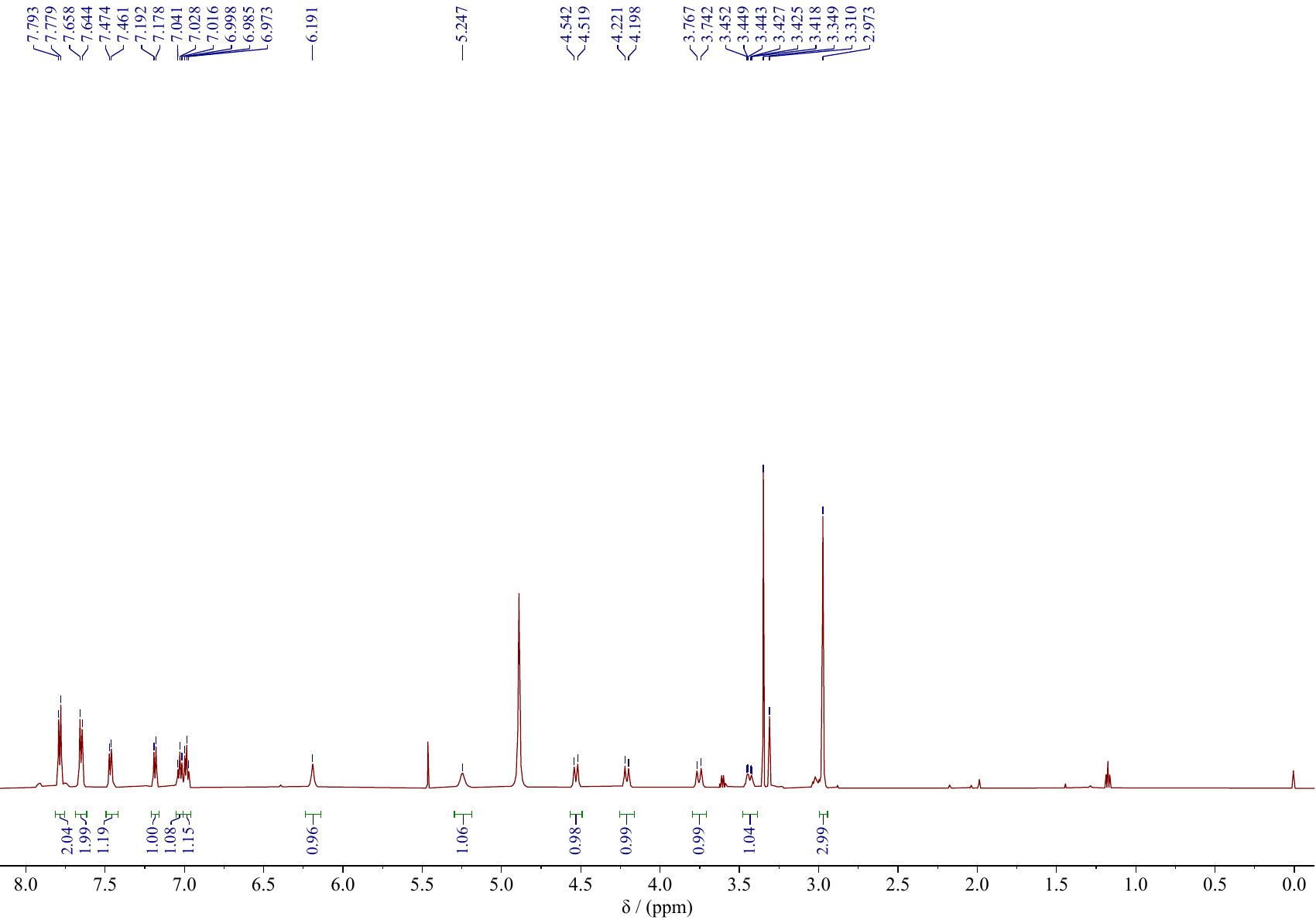

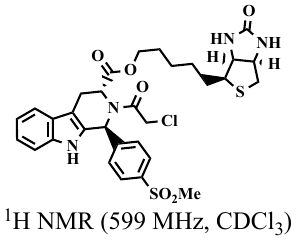

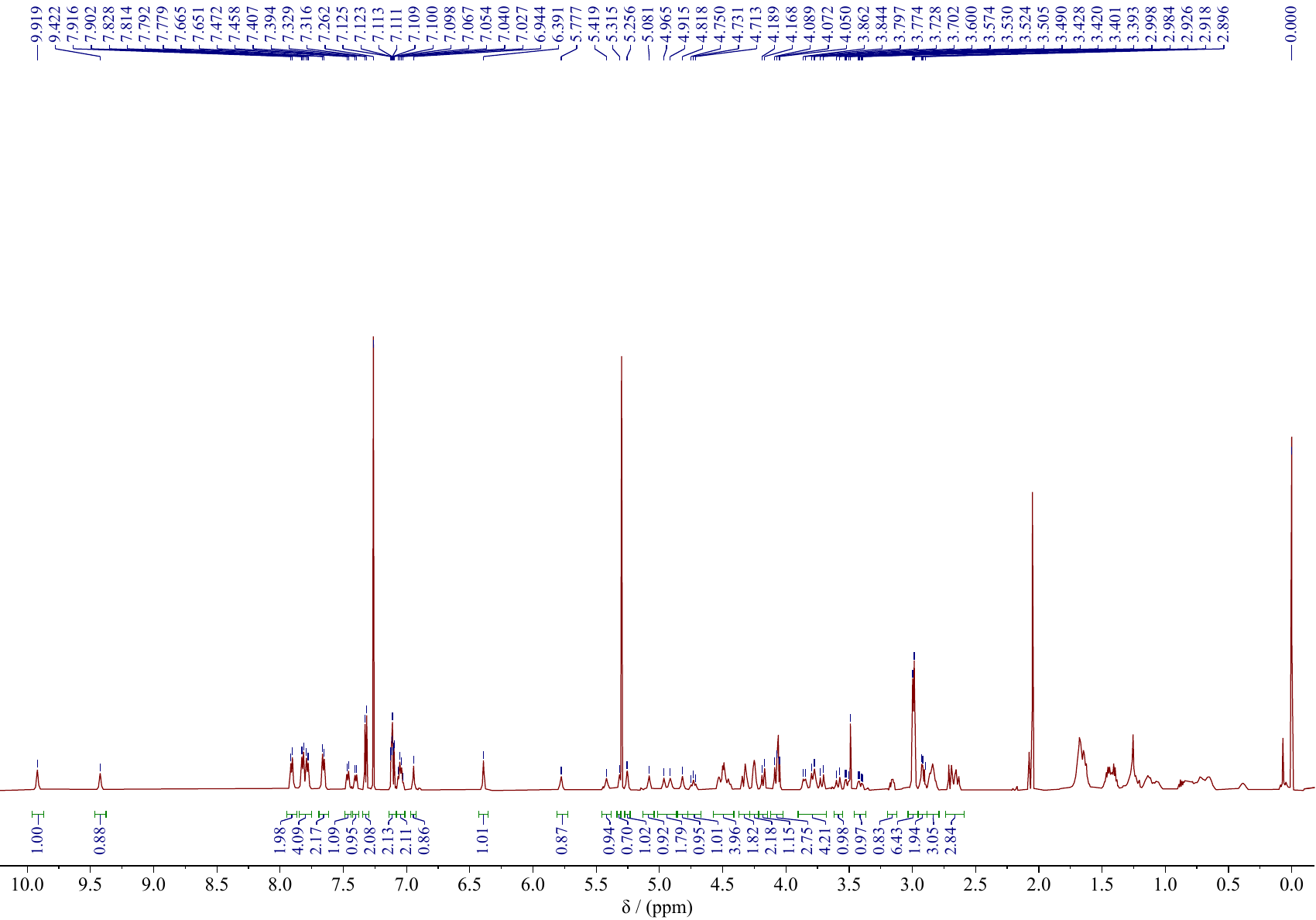

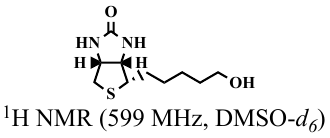

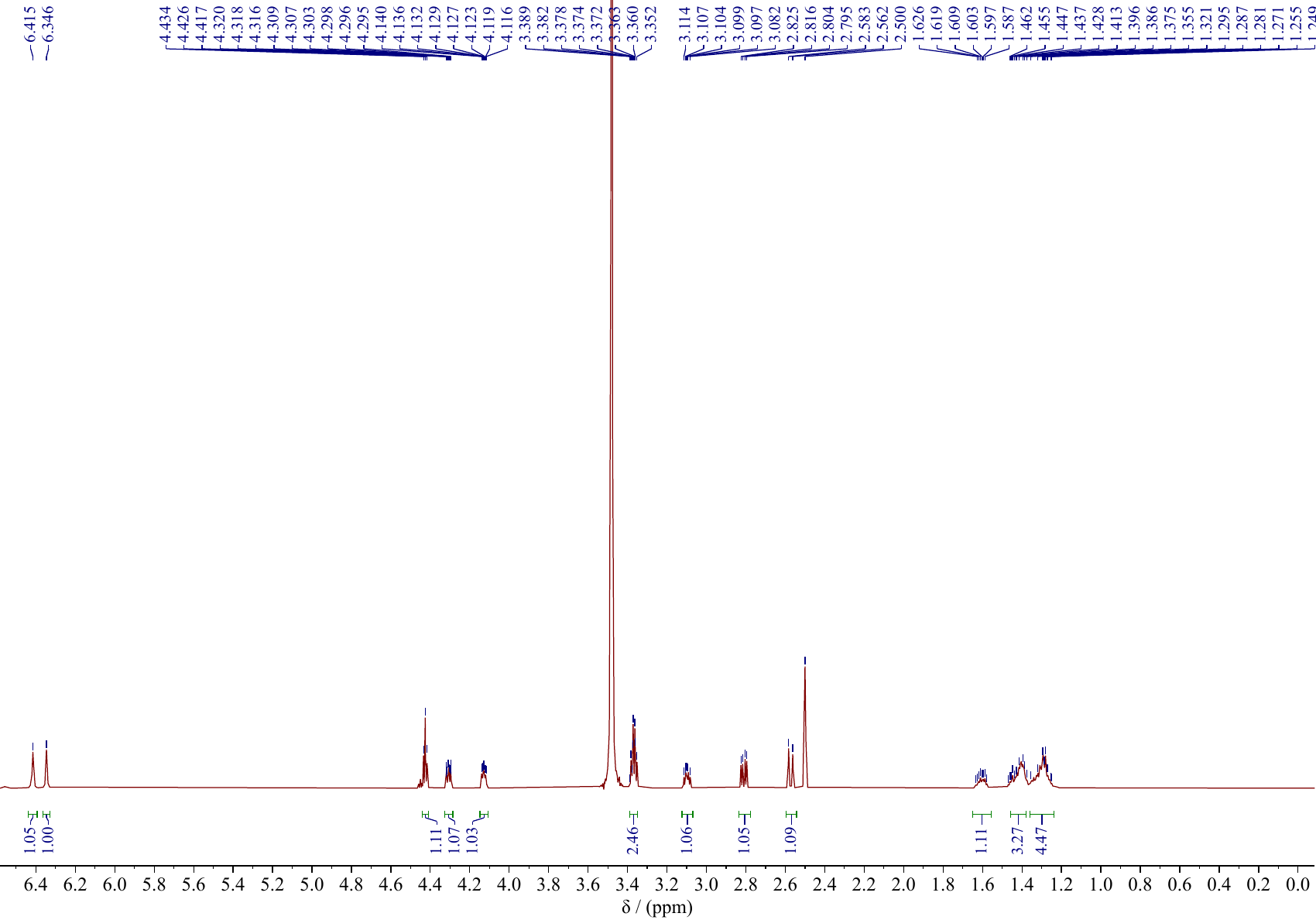

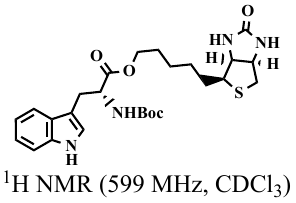

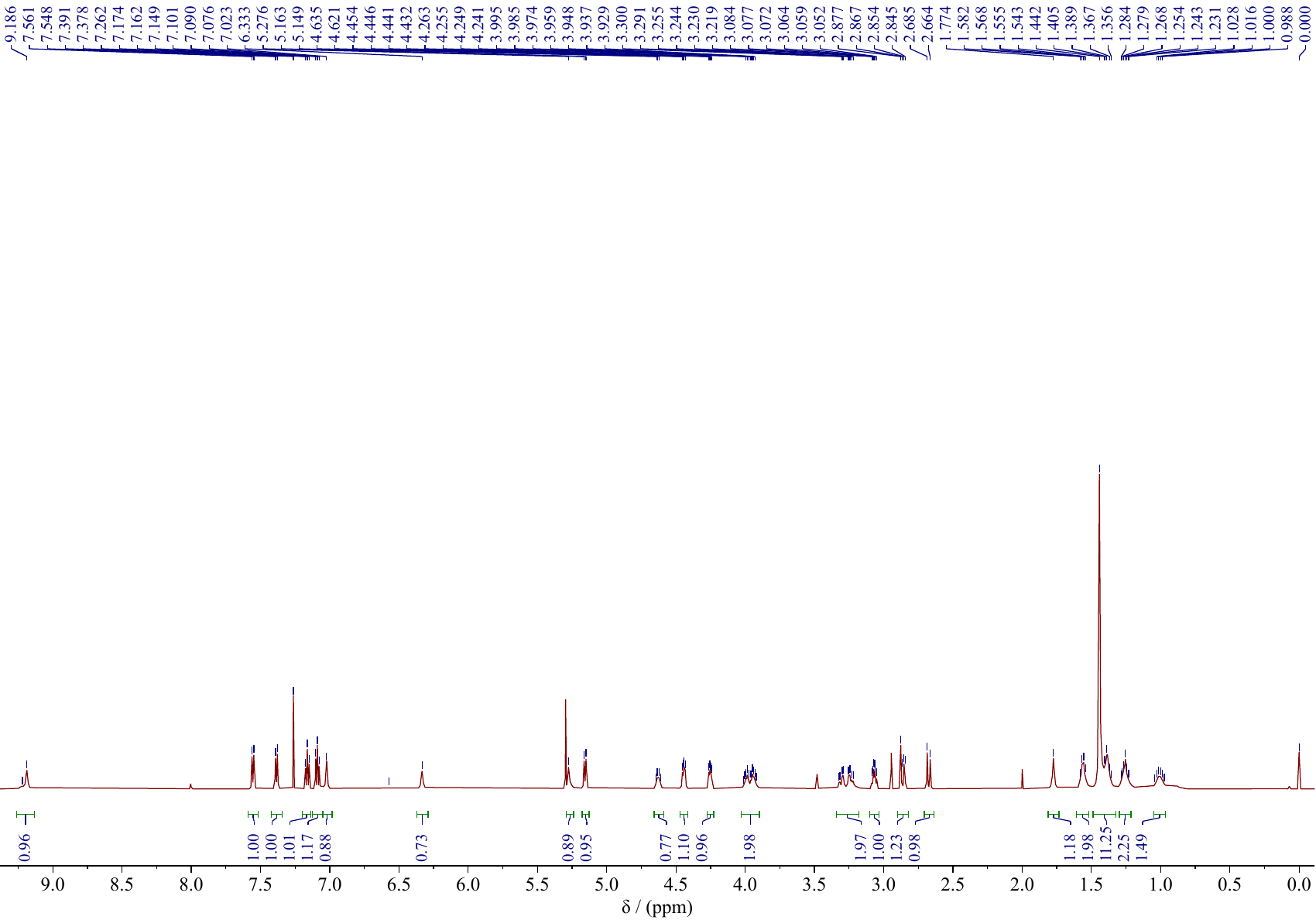

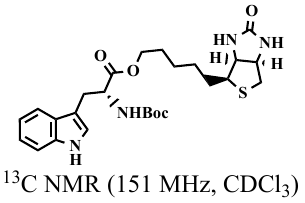

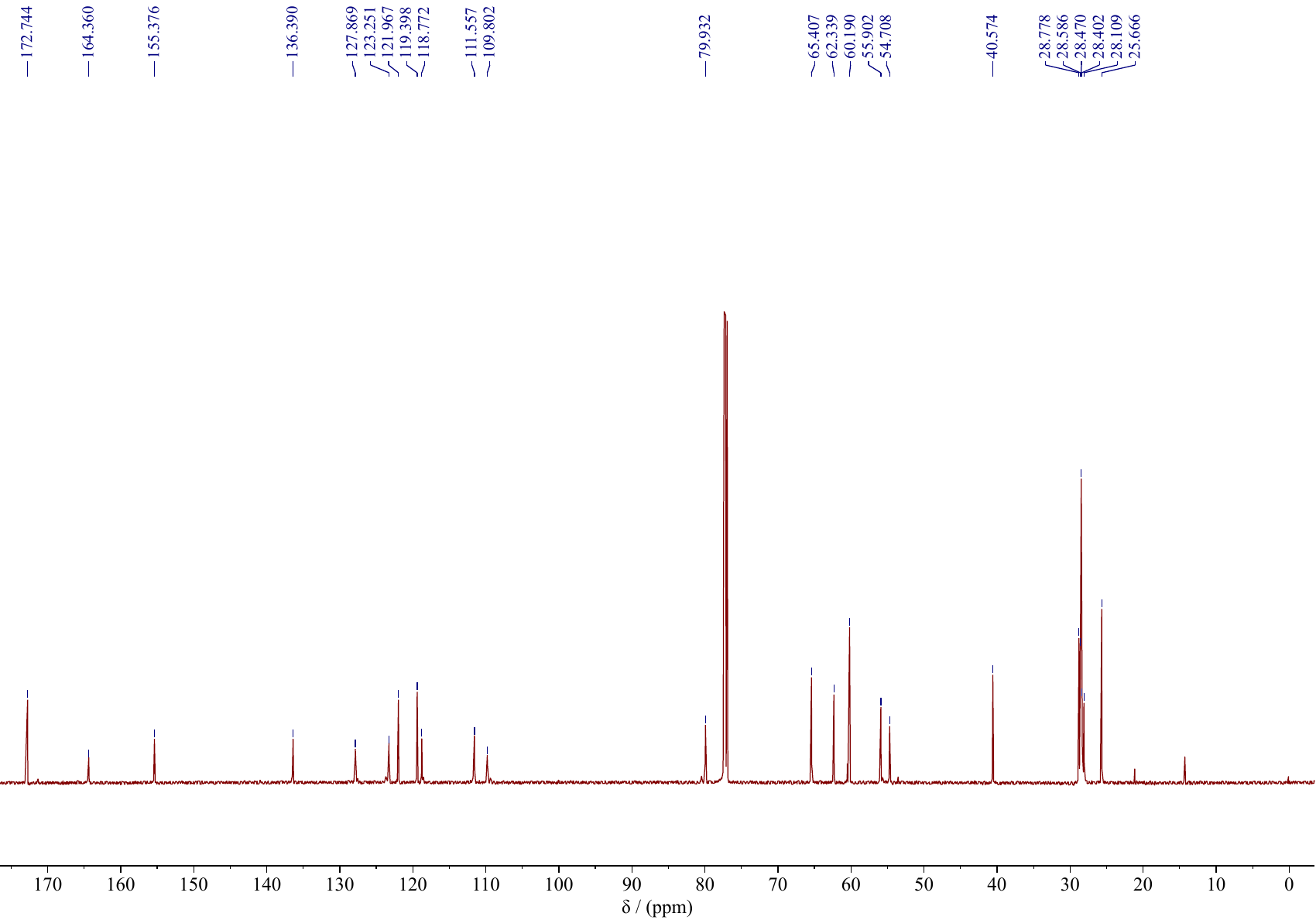

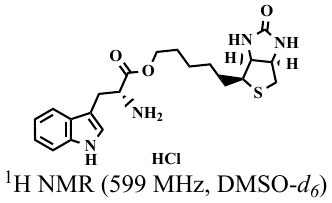

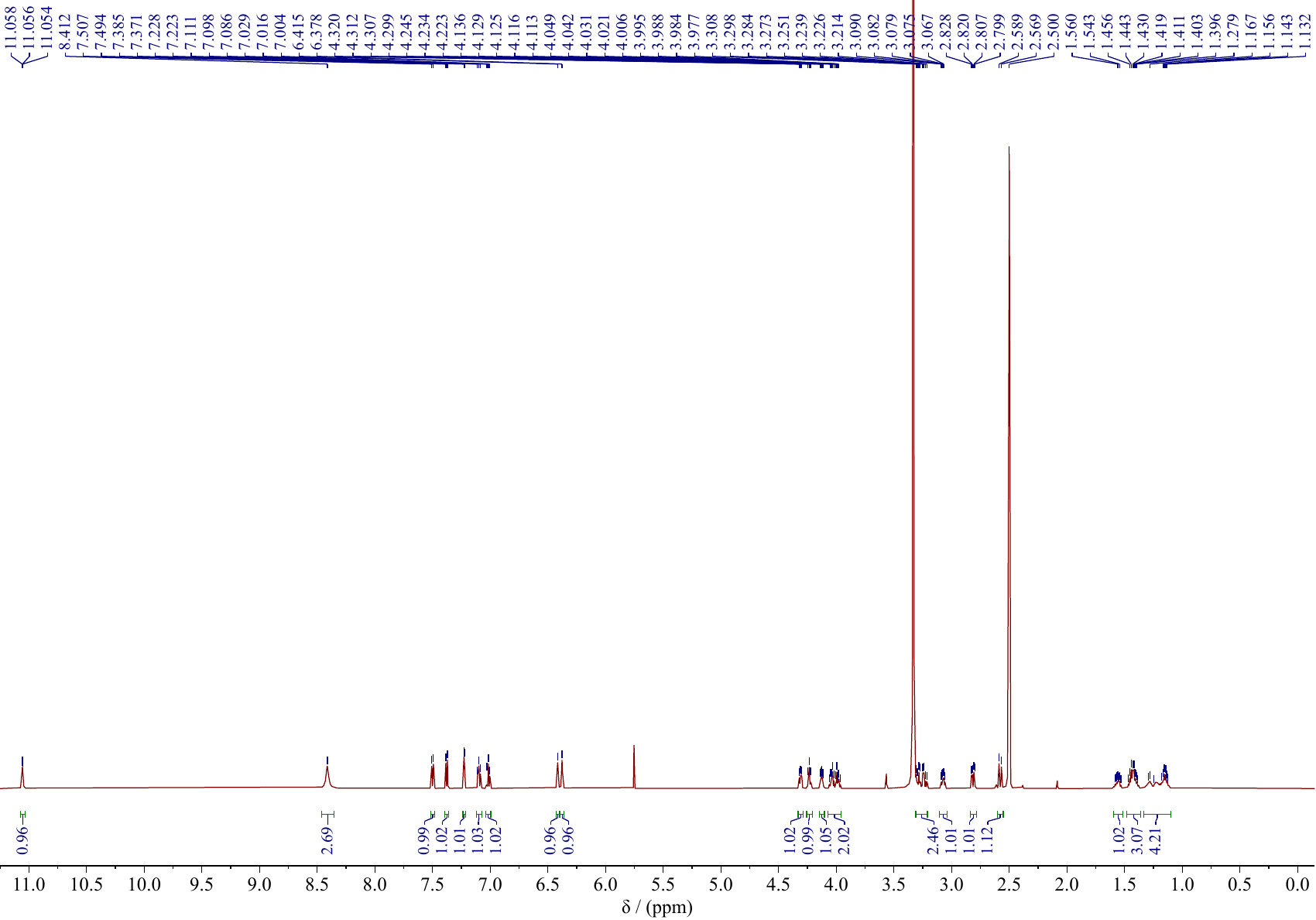

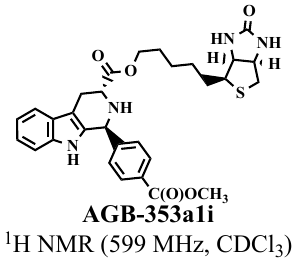

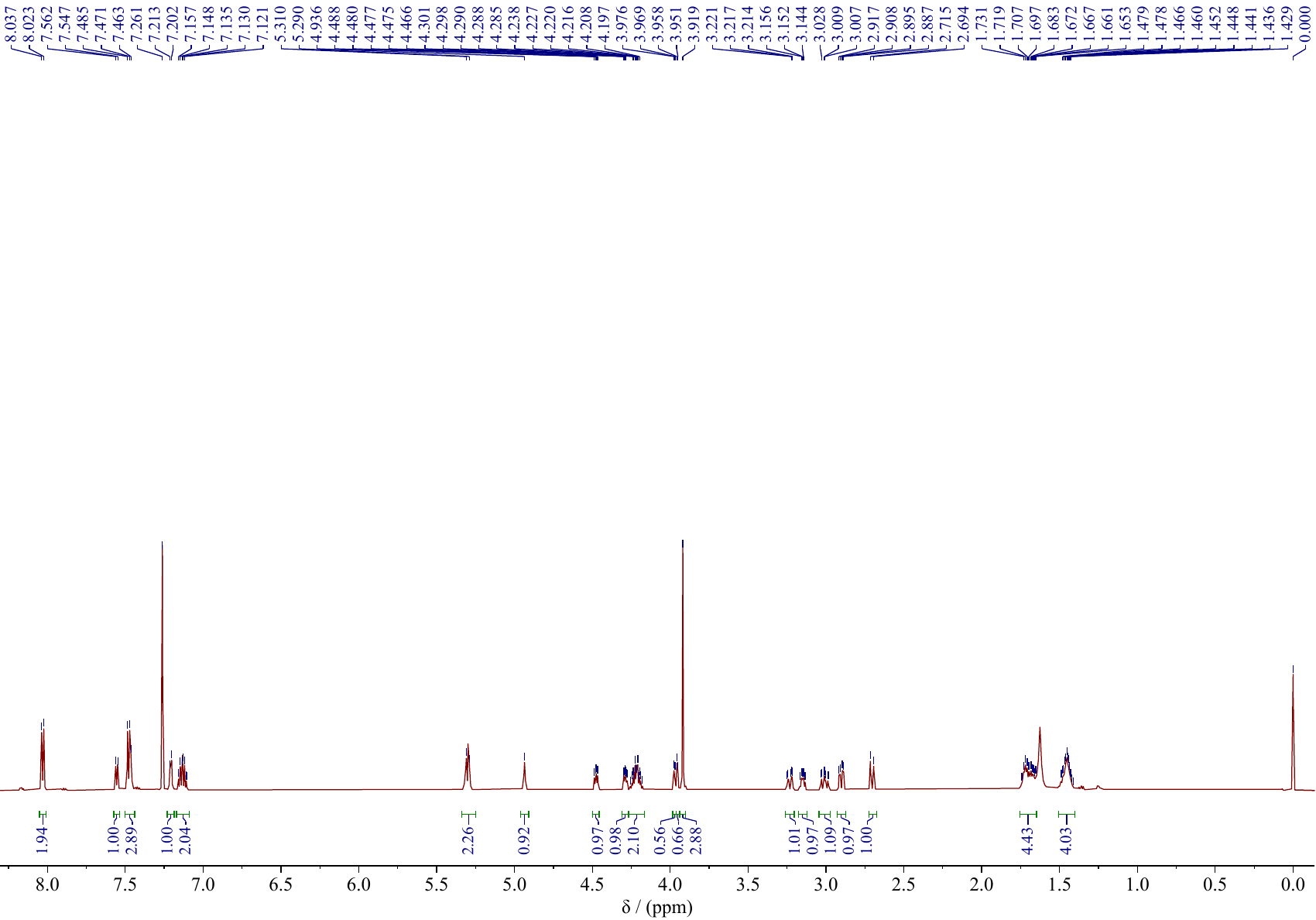

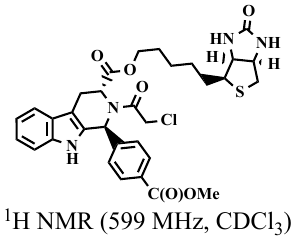

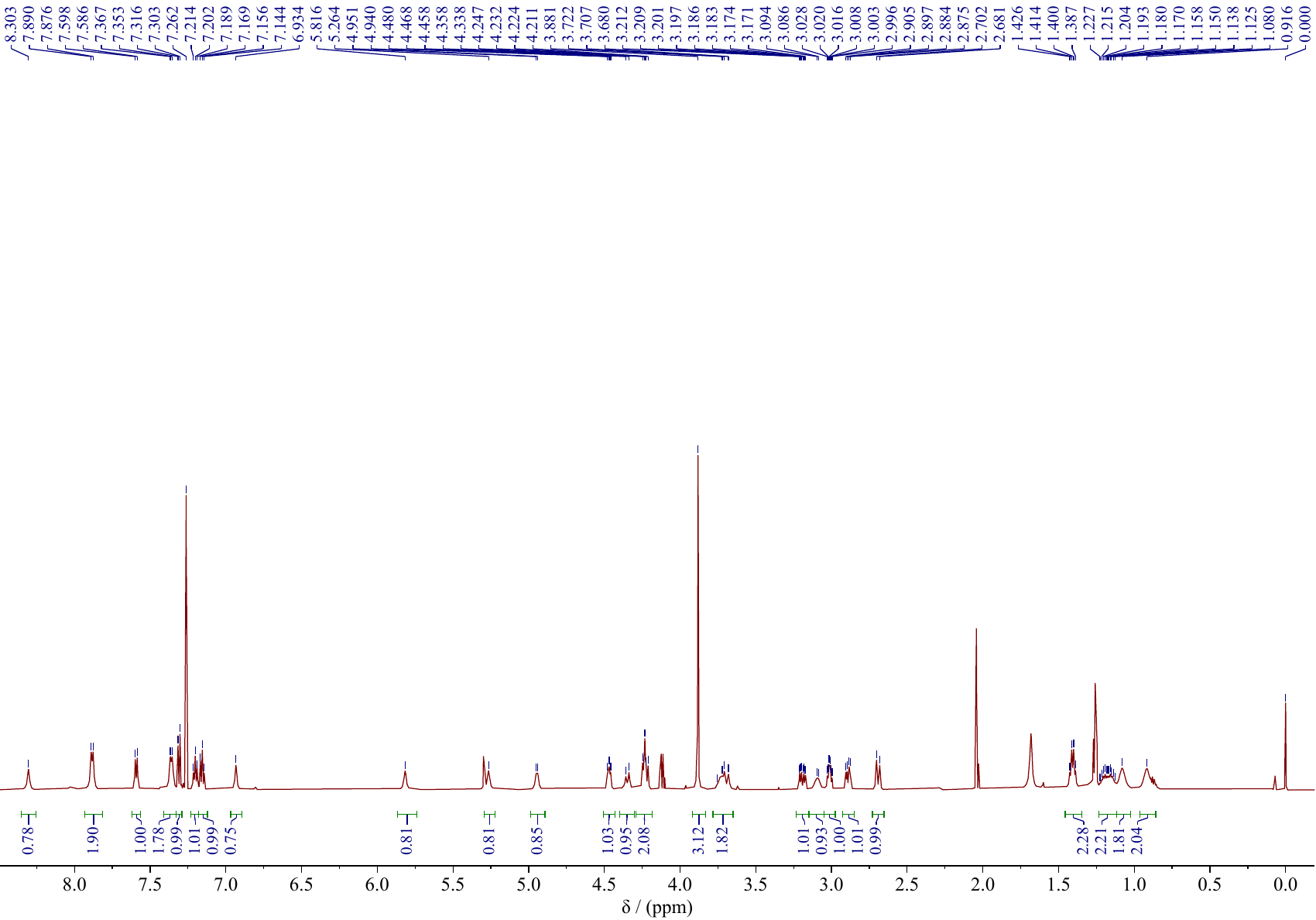

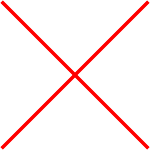

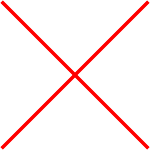

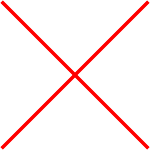

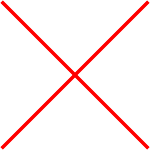

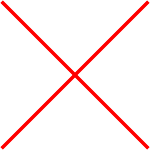

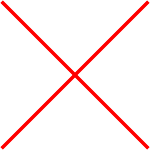

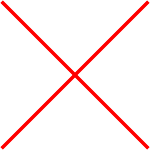

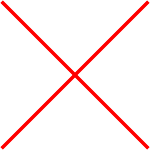

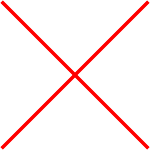

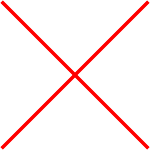

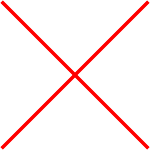
